# Efficient Cell Factory Design by Combining Meta-Heuristic Algorithm with Enzyme-Constrained Metabolic Models

**DOI:** 10.1101/2025.06.12.659423

**Authors:** Wenbin Liao, Gengrong Gao, Haoyu Wang, Xingcun Fan, Le Wang, Siwei He, Guangming Xiang, Xuefeng Yan, Hongzhong Lu

## Abstract

The rational design of high-performance microbial cell factories remains a central challenge in sustainable biomanufacturing due to the complexity of metabolic networks and the difficulty of predicting synergistic genetic interventions. Despite recent advances in strain design algorithms, predicting combinatorial targets remains computationally prohibitive due to the combinatorial explosion. Here, we present MetaStrain, a unified computational framework that integrates enzyme-constrained models (ecModels) with meta-heuristic algorithms to identify non-intuitive combinatorial gene targets for improving product yields. MetaStrain first performs pre-screening through a modified enforced objective flux scanning algorithm to reduce the dimensionality of candidate genes and annotate editing strategies. The subsequent search module employs flexible individual encodings compatible with diverse meta-heuristic algorithms and adopts a bottom-up phenotype evaluation strategy enabling efficient exploration of the combinatorial design space. Integrated redundancy analysis tools further identify single and fixed-size combinatorial strategies, facilitating direct experimental implementation. Computational simulation in *Saccharomyces cerevisiae* reveals significant enhancements in 2-phenylethanol and spermidine biosynthesis, while controlling target count and covering experimentally validated targets. Experimental validation in *Escherichia coli* further confirmed the algorithm’s predictive power, achieving up to a 61.25% increase in L-tryptophan titer of the five-target combination strain. Overall, MetaStrain achieves high computational efficiency, stable convergence, and broad adaptability across diverse metabolic targets. This study provides a powerful tool for metabolic engineering, bridging computational prediction and experimental realization, and highlighting the potential of meta-heuristic optimization in synthetic biology.

## 1. Introduction

In the 21st century, the rapid advancements in Artificial Intelligence (AI) and synthetic biology have significantly expanded the applications of microorganisms across diverse fields (Jullesson et al., 2015), including biofuels (Nielsen et al., 2013; Zhou et al., 2016), pharmaceuticals (Ro et al., 2006), food additives (Wei et al., 2018), and industrial enzymes (Duman-Özdamar and Binay, 2021). The increasing global demand for petroleum-based products, coupled with concerns about environmental sustainability, has driven a transition toward microbial fermentation-based bioproduction pathways as an alternative to traditional chemical synthesis (Nielsen and Keasling, 2016). However, succeeding in the competitive landscape of traditional chemical production requires the development of high-performance microbial cell factories (MCFs) (Volk et al., 2023). This necessitates the reprogramming of cells through metabolic engineering to create optimal strains with superior production metrics, including titer, rate, and yield (TRY) (Konzock and Nielsen, 2024).

Traditional strain optimization approaches rely heavily on researchers’ expertise, often guided by qualitative and intuitive design principles, requiring extensive trial-and-error experimentation. This process is both time-consuming, costly and limited in scope. To address these challenges, constraint-based modeling (Lewis et al., 2012) has facilitated the development of reliable quantitative metabolic models, providing a global perspective and a platform for system-level analysis in metabolic engineering. Genome-scale metabolic models (GEMs) provide a mathematical representation of metabolic networks, supporting pathway analysis, metabolic flux prediction, and strain design. GEMs can also integrate multi-omics data to improve predictive accuracy (O’Brien et al., 2015). To enhance model-based *in silico* strain design, the GEMs were further enhanced with enzymatic constraints using kinetic and omics data (Sanchez et al., 2017). This approach significantly reduces flux variability in traditional GEMs while revealing enzyme usage and distribution across metabolic pathways. It also overcomes the limitations of traditional GEMs in simulating metabolic shifts, such as overflow metabolism (Basan et al., 2015), the Crabtree effect (Nilsson and Nielsen, 2016), and the Warburg effect (Shlomi et al., 2011). The vast number of metabolites and reactions in these models results in high-dimensional solution spaces. Flux balance analysis (FBA) (Orth et al., 2010) is widely used to simulate metabolic phenotypes. Additionally, the minimization of metabolic adjustment (MOMA) (Segrè et al., 2002) was developed to simulate metabolic flux distribution in mutant strains following genetic interventions. Both FBA and MOMA have been extensively utilized for predicting gene targets in various computational strain design algorithms, but MOMA needs to be adjusted to ecModels.

A range of computational tools has been developed for *in silico* strain design, yet many faces significant limitations. Early methods, such as OptKnock (Burgard et al., 2003), employed bilevel optimization to identify gene deletion strategies, successfully increasing succinic acid production in *E. coli*. OptGene (Patil et al., 2005) pioneered the use of evolutionary algorithms for identifying deletion targets, optimizing production of compounds like succinic acid, glycerol, and vanillin in *S. cerevisiae*. However, these tools were limited to knockout strategies. To address overexpression targets, FSEOF (Choi et al., 2010) applies additional constraints on product formation fluxes, identifying genes with increased flux contributions, but lacks the ability to explore combinatorial editing strategies. More advanced tools, such as OptCouple (Jensen et al., 2019), extend the scope to include combinations of knockouts, insertions, and medium supplements, enabling growth-coupled production of target compounds. OptForce (Ranganathan et al., 2010) and OptDesign (Jiang et al., 2022) do not rely on alternative biological objectives, analyzing flux distribution differences to propose minimal intervention strategies. However, these tools largely rely on traditional GEMs, limiting their biological reliability. The ecFactory (Domenzain et al., 2025) successfully predicts engineering targets for 103 valuable chemicals using the ecModel of yeast, but it provides a large number of targets for each product. StrainOptimizer is a complete strain design toolbox based on Python, which further expands ecFactory to include the ETFL model (Wang et al., 2025). Incorporating advanced AI techniques, such as reinforcement learning and machine learning, into strain design has also been explored (Sabzevari et al., 2022; Zhang et al., 2020). Furthermore, these tools often struggle with the combinatorial explosion inherent in large-scale metabolic networks, making it difficult to identify optimal solutions (Lu et al., 2023).

Meta-heuristic algorithms, as representatives of symbolic AI, are powerful tools for solving complex combinatorial optimization problems due to their strong search capabilities and adaptability. Genetic algorithms (GAs) (Holland, 1992), for instance, simulate natural selection and heredity to achieve rapid searches in high-dimensional spaces. Differential evolution (DE) (Das and Suganthan, 2011), known for its simplicity and robustness, implement iterative updates through evolution operators, gradually approaching optimal solutions. Advanced strategies like elite retention and adaptive parameter control have significantly improved algorithmic performance (Deb et al., 2002; Huang et al., 2020). Meta-heuristic algorithms have shown great potential in addressing the complex optimization challenges of metabolic engineering, including high-dimensional and discontinuous solution spaces (Liu et al., 2025b; Wang et al., 2024; Yu et al., 2025). From pathway design to target prediction, phenotype simulation, and model parameter optimization, these algorithms provide effective tools for enhancing target product yields, optimizing strain performance, and uncovering metabolic network mechanisms. However, *in silico* strain design by combining Meta-heuristic algorithms and ecModels remains underexplored.

In this study, we present MetaStrain, a novel strain design framework that synergistically integrates ecModels with advanced meta-heuristic algorithms to identify non-intuitive combinatorial gene editing strategies for maximizing product yields in metabolic engineering. This integration enables MetaStrain to achieve high computational efficiency with stable convergence, facilitating the rapid and accurate identification of optimal gene editing strategies involving overexpression, knockdown, and knockout. The framework demonstrates strong adaptability and generalization, being compatible with all ecModels developed using the GECKO framework, which broadens its application across diverse species and metabolic objectives. Moreover, MetaStrain is modular and upgradable, supporting seamless incorporation of emerging search algorithms and gene-screening utilities to maintain scalability for future developments. MetaStrain successfully predicted diverse combinatorial targets that enhanced the biosynthesis of 2-phenylethanol (2-PE) and spermidine in *S. cerevisiae* while effectively controlling target number for experimental feasibility. Guided by predictions, combinatorial edits implemented in *E. coli* increased the L-tryptophan titer by up to 61.25%, confirming the accuracy and practical value of the framework. By addressing the computational challenges of large-scale combinatorial optimization and effectively guiding experimental design, MetaStrain represents a significant advancement in strain optimization, providing a powerful and innovative approach for reprogramming complex biochemical systems to achieve enhanced production metrics.

## 2. Materials and Methods

### 2.1 Framework of MetaStrain

The MetaStrain workflow (Fig. 1) comprises four key steps:

1. Model preprocessing and reference flux acquisition

**Fig. 1.**
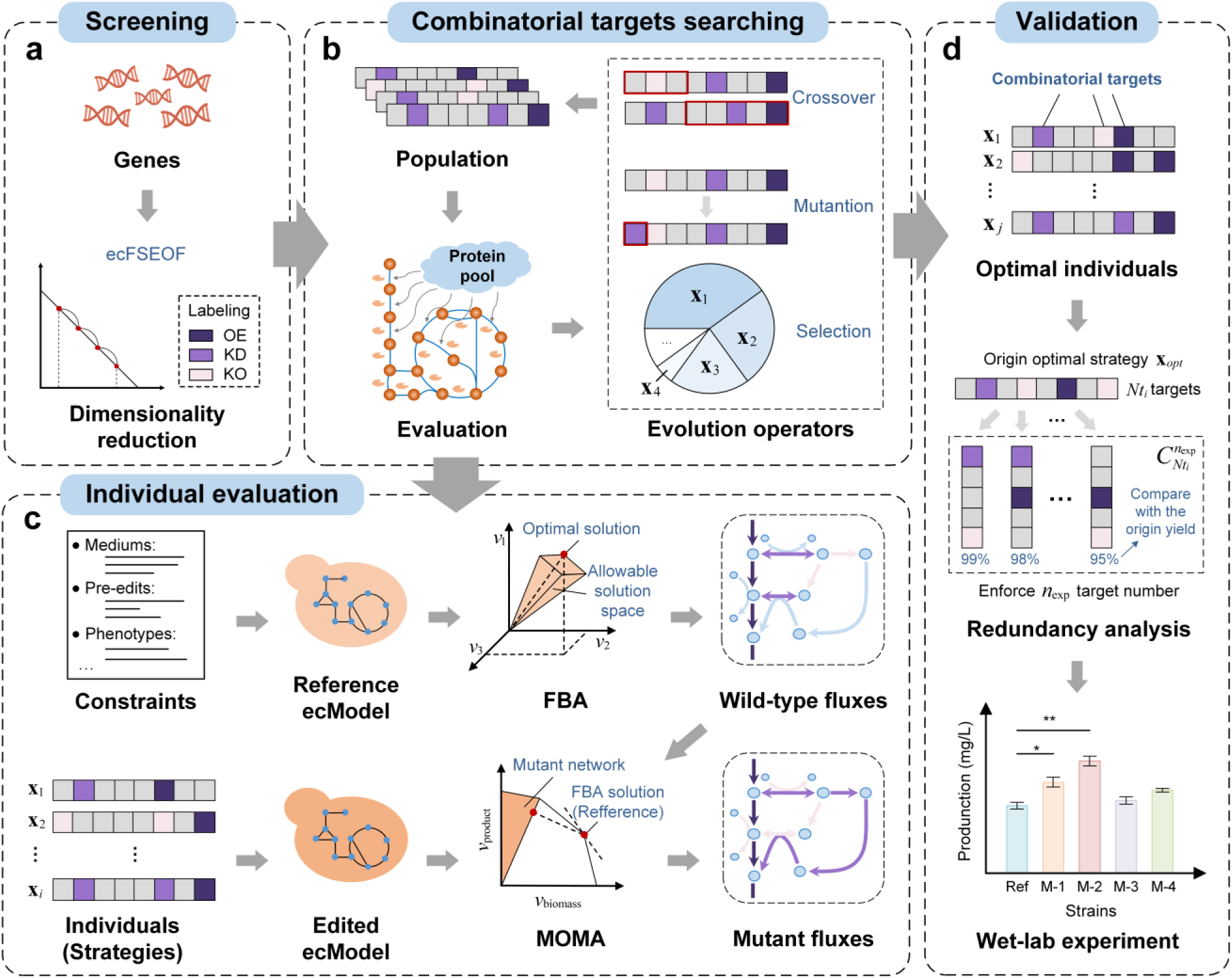
The computational framework of MetaStrain. The algorithm consists of two primary modules. (a) The initial screening module performs dimensionality reduction and annotation, where all enzyme-encoding genes in the metabolic model are systematically analyzed to pre-screen potential targets. (b, c) The search module, which executes the core optimization task. (b) The general iterative workflow of a meta-heuristic algorithm, where a population of candidate designs undergoes mutation, crossover, and selection. (c) The details of the fitness evaluation for each individual design. The process involves simulating a reference flux state and then solving for the flux distribution of the modified strain, ultimately yielding a fitness score for the search. (d) The wet-lab experiment verification. Statistical analysis and redundancy analysis were performed on the optimal individuals to obtain strategies with a fixed number of targets, which was feasible to verify in wet-lab experiments. The experimental verification provides a strong proof of the effectiveness of the algorithm.

For different microorganisms, ecModels are first adjusted and validated to accurately simulate metabolic states under different environmental conditions. This includes enabling oxygen, major carbon/nitrogen sources (with glucose as the sole carbon source in this study) and setting the upper bounds of reactions that initially defined as infinite to 1000. To enhance the resolution of protein synthesis-related fluxes, model precision is set to 10^−9^ . Essential genes and redundant metabolic pathways are removed. For exogenous products, relevant pathways are incorporated into the model, including adding new reactions, updating existing reactions, implementing gene manipulations, and setting constraints. The wild-type strain phenotype is then obtained through FBA by enforcing a minimum production level of the target product.

1. Gene dimension reduction and annotation

The ecFSEOF (Wang et al., 2025) method is used to perform gene dimension reduction and annotation for genes encoding enzymes in the model. To ensure that the randomly initialized individuals in a relatively small population can sufficiently cover the defined decision space, the number of candidate genes is reduced to fewer than 100. This significantly reduces the complexity of the search space, providing an efficient starting point for combinatorial target optimization.

1. Optimization process and algorithm implementation

MetaStrain integrates an efficient meta-heuristic optimization framework, using the yield of the target product as the objective function and encoding individuals according to the requirements of different algorithms. The main steps include population initialization, fitness evaluation, and evolutionary operations. Additional techniques, such as elite retention to accelerate convergence, regularization terms in the fitness function to control the number of targets, adaptive parameter control, and external archives, can further enhance algorithm performance.

1. Result analysis and validation

Due to the stochastic nature of the algorithm, ten independent experiments are conducted for each target product in a specific strain. The optimization results are scored using (Eq. (7)) and statistically analyzed to identify combinatorial targets with strong stability and biological significance. The results are further validated against experimentally verified targets comparation and pathway mechanism studies to assess their rationality and feasibility. In addition, each optimal strategy can be further analyzed for target redundancy, and an exhaustive search can yield fixed-size combinations, providing direct, actionable guidance for wet-lab implementation, for which experimental validation is essential to ensure the reliability of the predictions.

### 2.2 The ecModel of *S. cerevisiae*

GEMs are widely used to represent the metabolic networks of microorganisms by integrating genes, enzymes, and reactions through gene-protein-reaction (GPR) rules. These models encode microbial metabolism in a *M* × *N* dimensional stoichiometric matrix (**S** matrix), where rows represent metabolites, columns correspond to reactions, and matrix elements denote stoichiometric coefficients. The GECKO method expands GEMs by introducing additional rows and columns into the **S** matrix to explicitly include enzymes as reaction components. This approach highlights a fundamental constraint on metabolic fluxes (1): each flux (*v _j_*) cannot exceed the maximum enzymatic rate (*v*_max_), which is determined by the intracellular concentration of the enzyme and its turnover number.

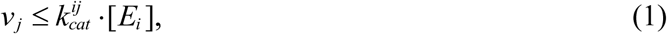

Here, [*E_i_*] denotes the intracellular concentration of each enzyme *E_i_*, and 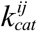 is the turnover rate. This quantitative method imposes soft constraints at the enzyme level, integrating proteomics data. In cases where detailed proteomics data are unavailable, a global enzyme mass constraint, similar to the FBAwMC (Beg et al., 2007) method, is applied. This introduces a pseudo-metabolite termed the "protein pool", to ensure that all metabolic reaction fluxes remain within biologically reasonable limits. For this study, we used the ecModel of *S. cerevisiae* ecYeast8.3.4, a well-characterized industrial chassis for metabolic engineering. Developed and refined by Lu et al. (Lu et al., 2019) (Table S1).

### 2.3 The ecModel of *E. coli*

*E. coli* is a cornerstone model organism in metabolic engineering, featuring a well-characterized genetic background, rapid growth, and mature, reliable genome-editing tools. High-quality GEMs for *E. coli* are available (Monk et al., 2017). Building on these, Ye et al. integrated enzymatic kinetic information to construct the enzyme-constrained model eciML1515 (Table S1) (Ye et al., 2020). To ensure close agreement between computational predictions and our experimental system, we introduced strain- and condition-specific constraints on top of eciML1515 and performed targeted model refinement.

Specifically, the baseline L-tryptophan-producing strain MGHT-10-1 carries knockouts in b2600, b3708, and b2599, and overexpresses terminal L-tryptophan biosynthesis genes b1260, b1261, b1262, b1263, and b1264. In the model, reactions corresponding to knockouts were constrained by setting their upper bounds to 0. For overexpression, due to the lack of directly calibratable reference fluxes or fold-change measurements, we adopted a phenotype-alignment strategy: we fixed the model’s L-tryptophan synthesis rate and glucose uptake rate to the experimentally observed values of 0.02 mmol gDWh^-1^ and 1.8 mmol gDWh^-1^, respectively, and performed FBA to maximize biomass. This constraint set substantially elevates flux through the relevant pathways compared to the unaligned model, thereby providing an effective model-level representation of overexpression.

Regarding medium and environmental constraints, we matched the experimental medium formulation by removing unavailable ions from the model (e.g., Fe³⁺, Ni²⁺, and MoO₄²⁻) and adding accessible nutrients derived from yeast extract, including tyrosine, phenylalanine, and citrate. During the algorithmic search phase, to maintain feasibility and avoid non-physiological extreme solutions, we imposed only minimal lower bounds on growth and product formation—specifically, a minimum growth rate of 0.1 and a minimum product formation rate of 0.01—while leaving other bounds at the model defaults.

### 2.4 Phenotype simulation of wild-type and mutant strains

#### Flux balance analysis

FBA employs linear programming techniques, such as the simplex method, to solve optimization problems. FBA predicts the steady-state metabolic phenotype of a microorganism under specified environmental conditions, without relying on complex kinetic parameters. For single-celled organisms, this objective often aligns with maximizing biomass synthesis, under the assumption that these organisms have evolved to achieve optimal growth performance.

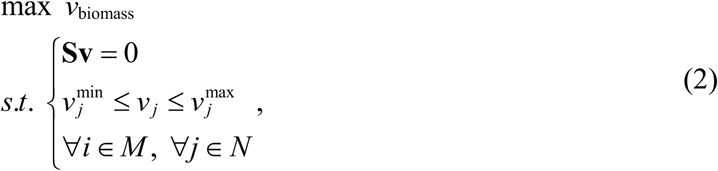

Here, *v*_biomass_ represents the flux of biomass production. **Sv** = 0 provides a global equality constraint for the model, ensuring mass conservation. 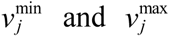 represent the minimum and maximum bounds of a reaction *v _j_* . The problem can be solved using a linear programming solver, such as Gurobi (Gurobi Optimization).

In this study, the wild-type strain’s metabolic phenotype serves as the baseline for subsequent computational analyses, which was predicted by a single-step FBA-based approach. First, the flux range of product was first set approximately half of its theoretical maximum flux, ensuring that the simulated strain maintains a certain production level. Biomass synthesis was then maximized as the objective function for solving.

### Phase plane analysis

Phase plane analysis can be performed by gradually varying the target flux and constraint conditions. It provides a comprehensive understanding of the global characteristics of metabolic networks, particularly trade-off relationships between two or more metabolic fluxes. In this study, the StrainDesign (Schneider et al., 2022) toolbox was used to construct phase planes for a target product, biomass synthesis, and glucose consumption. This method allows for quick and intuitive insights into nonlinear couplings between fluxes of interest, identifying the factors influencing product yields.

### Minimization of metabolic adjustment in ecModel

Genetically engineered strains often exhibit suboptimal metabolic flux distributions due to the absence of natural evolutionary processes, with an initial tendency to resemble the wild-type flux. To predict these suboptimal states, the MOMA method minimizes the Euclidean distance (Eq. (3)) between the flux profiles of wild-type and mutant strains, formulated as a quadratic programming problem. While effective for standard GEMs, MOMA requires adaptation for ecModels due to their unique structure. Here, we introduce an improved method, ecMOMA.

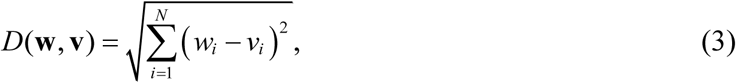

Here, **w** represents the metabolic flux vector of the wild-type strain, serving as the reference vector in the algorithm, and **v** represents the flux distribution of the mutant strain.

In ecModels, each enzyme is associated with a “synthesis reaction” and an “arm reaction” to ensure total flux constraints. However, the flux magnitudes of enzyme-synthesis reactions significantly lower than those of the original metabolic reactions. This discrepancy causes MOMA results to be dominated by the original metabolic reactions, and the introduced arm reactions become redundant, leading to double-counting of certain metabolic fluxes. To resolve this, ecMOMA refines the distance function by excluding enzyme-synthesis reactions, arm reactions, and protein pool exchange reactions from the flux space. This adaptation improves simulation accuracy, reduces computational redundancy, and enhances efficiency in both time and memory usage.

### 2.5 Gene dimension reduction and labeling using ecFSEOF

The ecFSEOF algorithm (Domenzain et al., 2025) was employed to identify gene editing targets. This method integrates the features of GECKO, extends its application to ecModels, and incorporates a scoring function to prioritize genes for overexpression, knockdown, or knockout. In this study, the algorithm sets the biomass synthesis flux to its theoretical maximum and progressively decreases it across predefined levels to explore metabolic changes. Flux values are obtained through FBA with the target product yield as the objective function, followed by a two-step scoring process to identify potential engineering targets. For a detailed description of the scoring methods and criteria for gene prioritization, please refer to the supplementary materials (Supplemental Material).

### 2.6 Characterization of gene editing in ecModels

In MCF design, representing gene editing operations—specifically, overexpression, knockdown, and knockout of genes, and determining the extent of these edits—remains challenging. The ecModels address this by modulating enzyme usage, as they treat enzymes as key components directly involved in metabolic reactions. Enzyme synthesis reactions establish a one-to-one correspondence between enzymes and genes, while other reactions represent enzyme utilization. These relationships are embedded in the metabolic network and adhere to mass conservation principles.

In ecYeast, gene knockout was implemented by setting the flux bounds of the corresponding enzyme synthesis reaction to zero, effectively deactivating the pathway. For overexpression and knockdown, flux bounds were manually adjusted based on reference enzyme usage values derived from the wild-type phenotype. Overexpression was defined as increasing the lower flux bound to four times its reference value, while knockdown reduced the upper flux bound to half of its reference value. If the reference flux value is 0, then the knockdown and knockout operations represented in the model are the same. We uniformly regard the knockdown-targets as knockout-targets, indicating that the relevant reactions in the mutant strain are also not allowed to have flux. These adjustments were informed by literature and practical considerations to ensure rational and effective interventions (Segrè et al., 2002). To address the issue of zero flux predictions for many reactions due to model precision limitations, a minimum flux value (*ε* = 4×10^−9^) was introduced. For genes targeted for overexpression with an initial flux of zero, the lower flux bound was set to 4*ε*, ensuring feasibility within the model’s precision constraint.

### 2.7 Meta-heuristic algorithm

Meta-heuristic algorithms operate through iterative processes and parallel searches to identify satisfactory solutions, rather than absolute optimal. These algorithms are especially suited for global optimization problems, as they balance exploration and exploitation. While GA and DE share similar iterative processes, they differ in their evolutionary operators and auxiliary search strategies. In this study, GA and adaptive differential evolution with optional external archives (JADE) (Zhang and Sanderson, 2009) were employed for combinatorial target searches. GA was divided into two forms, binary-coded (binGA) and symbolic-coded (symGA), based on whether gene annotations were available. While JADE’s adaptive parameter control and external archives increase computational overhead, they offer superior global optimization capabilities and can achieve effective results without relying on gene annotations.

Innovatively, we designed parameters within GA and JADE initializations to ensure population uniformity and individual feasibility. Appropriate thresholds maintained a low number of targets, ensuring experimental economic viability. Individual encoding complied with algorithm requirements; for example, JADE operated within a continuous space. These initialization strategies avoid excessive interventions, thus promoting feasible and economically viable solutions. In binGA and symGA, a roulette wheel selection strategy was used to select candidate individuals, and an elite retention strategy ensured the direct inheritance of top performers to the next generation. Threshold control during single-point crossover and mutation further optimized population updates (Supplemental Material). JADE introduces unique mutation strategies DE/current-to-*p*best*s*, optional external archive and adaptive parameter control to improve its global optimization capabilities. The external archive adds diversity during iterations, ensuring effective search direction. Detailed JADE operating procedures and equations are available in the supplemental material.

### 2.8 Evaluation of individuals

In meta-heuristic algorithms, the quality of each individual is assessed through its fitness, with higher fitness values indicating better solutions. TRY are valuable economic indicators in fermentation processes. In this study, the yield of the target product is selected as the objective function, defined as:

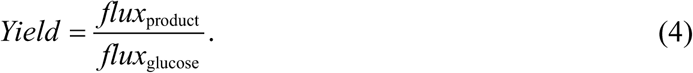

During the evaluation process, the wild-type strain phenotype is first obtained as a reference flux through FBA of the original model. Then, the ecModel is modified according to the gene editing strategy represented by the individual. These two inputs are used in ecMOMA to compute the mutant strain phenotype and subsequently calculate the target product yield. Notably, a minimum growth capacity is ensured for mutant strains, with biomass synthesis flux constrained to no less than 0.1 mmol gDWh^-1^ . This bottom-up design makes MetaStrain inherently suited for identifying synergistic combinations of targets.

We desinged two types of fitness function. First, we directly took yield as the fitness value (Eq. (5)). Then we consider adding a regularization term to the function to control the number of combined targets (Eq. (6)).

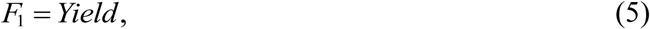

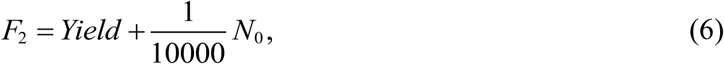

Here, denotes the number of dimensions encoded as 0 in the individual, and is scaled to adjust the influence of the target count on the fitness value. Other regularization terms, like the double hook function, show the same result as the linear term while requiring additional computing resources.

It is important to note that the ecMOMA calculation is essentially a quadratic programming problem, and models with extensive gene edits may not always yield optimal solutions. The solver may return statuses such as “optimal”, “infeasible” or “unbounded”. To ensure the algorithm proceeds smoothly, non-optimal cases are assigned a small nonzero fitness value (0.001). This assignment ensures that these individuals have a low probability of selection without significantly affecting population quality, thereby increasing the algorithm’s diversity.

### 2.9 Ranking of optimal individuals

Through repeated experiments, we can obtain the optimal editing strategies found in each search, along with their corresponding fitness values. Following the search, strategies were ranked based on three combined criteria: the regularized fitness values, the total number of targets, and the number of experimentally validated targets. By applying weighted summation to these components, we derived a prioritization strategy to guide subsequent wet lab experiments.

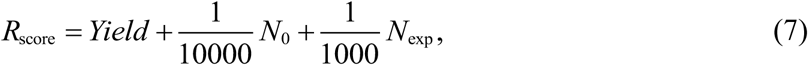

where *N*_exp_ represents the number of experimentally validated targets in the strategy. This comprehensive ranking approach ensures that the selected gene editing strategies not only maximize yield but also align with experimental validation results.

### 2.10 Eliminate redundant analysis

Meta-heuristic algorithms naturally account for combinatorial interactions among targets. However, when using fitness function *F*_1_, the number of targets does not directly influence fitness, which can admit redundant edits that weakly relate to the product and consume limited resources without changing the objective. When using *F*_2_, marginal cases may arise where the benefit of adding a small number of targets is offset by the penalty term. To obtain more economical, implementable, and interpretable designs, we conduct a post hoc redundancy analysis on the elite set returned by the algorithm.

### Analysis of single-target redundancy

For each optimal editing strategy *x _j_*, containing *n* targets, we iteratively evaluate the set of *n* derived strategies formed by removing one target at a time, each with (*n* −1) targets. If removing target *i* reduces fitness to below 95% of the original strategy’s fitness, we classify target *i* as “essential”; otherwise, it is classified as “redundant.” Notably, for certain target groups, removing any single member may not reduce fitness, whereas removing multiple simultaneously can markedly degrade performance—precisely reflecting MetaStrain’s ability to capture higher-order combinatorial effects.

### Screening for fixed-size combinatorial strategies

Considering experimental timelines, construction complexity, and resource constraints, practical implementations often favor a fixed number of interventions *m* (typically 3–5). Accordingly, we enumerate all size- *m* subsets from the target set of each optimal or elite strategy and perform *in silico* evaluation and ranking of these combinations to generate a directly actionable candidate list for experiments. Because the total number of targets in the algorithm’s optimal individuals is far smaller than the entire candidate pool, this enumeration can be performed efficiently.

### 2.11 Strains and plasmids

All strains and plasmids used in this study are listed in Table S2 and Fig. S1 *E. coli* DH5α was employed for plasmid construction and transformation. Genome editing utilized a two-plasmid CRISPR–Cas9 setup, with pEcCas driving Cas9 expression via an arabinose-inducible promoter and pEcgRNA supplying the target-specific sgRNA (Li et al., 2021).

### 2.12 Plasmids and strains construction

pEcgRNA-derived constructs were assembled by Gibson assembly. Upstream and downstream homology arms (∼500 bp each) were fused using overlap-extension PCR. Target-gene modules were amplified with KOD One High-Fidelity DNA Polymerase (TOYOBO, Japan) and joined by seamless multi-fragment cloning (Vazyme, Nanjing, China). The pEcCas plasmid, pEcgRNA plasmid, and a donor fragment bearing the target gene were co-introduced into electrocompetent *E. coli* by electroporation. Transformants were selected on spectinomycin (50 μg/mL) and kanamycin (50 μg/mL). Plasmids were then cured sequentially: pEcgRNA by growth in LB containing 10 mM rhamnose, and pEcCas by counterselection in LB containing 1% sucrose. All primers are provided in Table S3.

### 2.13 Shake-flask fermentation

Shake-flask fermentations of *E. coli* MGHT-10-1 and derivative strains were conducted using a two-stage cultivation scheme optimized for metabolic engineering. Seed cultures (15 mL) were grown in 250 mL shake flasks at 37 °C and 220 rpm for 12 h; the seed medium formulation followed previous report (Xiong et al., 2021). Log-phase seed (2.5 mL; 10% v/v inoculum) was then transferred into 22.5 mL of fermentation medium (25 mL working volume per 250 mL flask) and cultivated for 48 h in fed-batch mode under the same temperature and agitation. The fermentation medium composition matched that described previously (Xiong et al., 2021). To enable real-time, colorimetric pH tracking, phenol red was included at 2% (w/v), and the culture pH was maintained near neutrality by titration with ammonium hydroxide (28%, v/v).

### 2.14 Analytical methods for quantifying L-tryptophan

L-tryptophan was quantified by high-performance liquid chromatography (HPLC; Agilent 1220, Agilent Technologies, USA). Separation was achieved on an Agilent Eclipse AAA column (4.6 × 150 mm, 3.5 µm) held at 39°C, with UV detection at 278 nm. Isocratic elution employed 0.3 g/L KH_2_PO_4_ (solvent A) and methanol (solvent B) mixed at 90:10 (A:B), delivered at 1.0 ml min^−1^ (Shen et al., 2012).

## 3. Results

### 3.1 MetaStrain: a novel framework for optimizing metabolic engineering strategies

The MetaStrain framework is engineered to design optimal metabolic engineering strategies through a modular computational pipeline. This species-independent framework consists of two primary modules (Fig. 1a, b): the initial screening module, and the search module. Benefiting from its modular architecture, MetaStrain can incorporate various strain design algorithms for dimensionality reduction, thereby complementing and extending their applicability. Moreover, it allows the integration of advanced optimization algorithms from the field of intelligent computation. Together with our customized encoding scheme and unique evaluation process, these components enable fully automated prediction of combinatorial target strategies across the entire workflow.

In the first module, we adopt the first step of our previous algorithm, ecFactory (Domenzain et al., 2025), to reduce the dimensionality of enzyme-encoding genes, thereby generating a prioritized candidate set for downstream optimization. In the search module, each individual encodes a combinatorial gene editing strategies and ecMOMA (Materials and Methods) is used to simulate the phenotype of the corresponding engineered strains (Fig. 1c), with target product yield (Eq. (4)) serving as the fitness value. For rapid prediction, we first employ a binary-coded GA (binGA) that relies on both dimensionality reduction and target annotation from the screening module. In addition, two alternative meta-heuristic algorithms—a symbolic-coded GA (symGA) and adaptive differential evolution with optional external archives (JADE)—can be independently applied to the reduced gene list. Unlike binGA, these two algorithms autonomously determine the manipulation type (overexpression, knockdown, or knockout) for each gene, without relying on prior annotation. Although more computationally intensive, this approach eliminates potential biases from the initial screening, broadens the exploration space for target interactions, and remains compatible with unannotated dimension-reduction workflows. During the search, an elite retention strategy can be incorporated to accelerate convergence, while the number of gene targets selected for edition could be controlled through the optimization of fitness function or parameters (Materials and Methods).

To enhance robustness of our framework, we conducted multiple independent optimization runs. This enabled statistical analysis of the results and allowed us to select the most promising strategy based on a comprehensive criterion (Eq. (7)). The optimized outcomes were further refined by identifying and removing redundant targets, which facilitated the enumeration of the optimal combinations of gene targets with a fixed size that maintain a balance between predicted performance and experimental feasibility. The versatility and efficacy of MetaStrain were demonstrated through case studies on designing MCFs for various target products and host strains. Some of these results were further subsequently validated through wet-lab experiments, as detailed in the following sections.

### 3.2 *In silico* evaluation of MetaStrain for gene target prediction to enhance 2-Phenylethanol production in *S. cerevisiae*

We first evaluated MetaStrain on the production of 2-phenylethanol (2-PE), which is a high-value aromatic compound used in food, cosmetics, and pharmaceuticals (Wang et al., 2019), to benchmark its ability to navigate combinatorial design spaces and recover biologically meaningful strategies in *S. cerevisiae*. Three-dimensional phase plane analysis revealed significant metabolic trade-offs between 2-PE production, biomass synthesis, and glucose consumption (Fig. S2a), underscoring the need for rational adjustments to the metabolic network to optimize resource allocation without compromising cellular viability. Simulation using the ecYeast8.3.4 (Lu et al., 2019) indicated a theoretical maximum flux of 4.9009 mmol gDWh^-1^ for 2-PE production, with an initial fitness value of 0.1231. Based on ecFSEOF (Wang et al., 2025), we identified 52 candidate genes (Table 1), each with a specified editing direction : overexpression, knockout, or knockdown. We further designed flexible individual encodings (Fig. 2a) that can interface with various algorithms, demonstrating the versatility of the proposed framework.

**Fig. 2.**
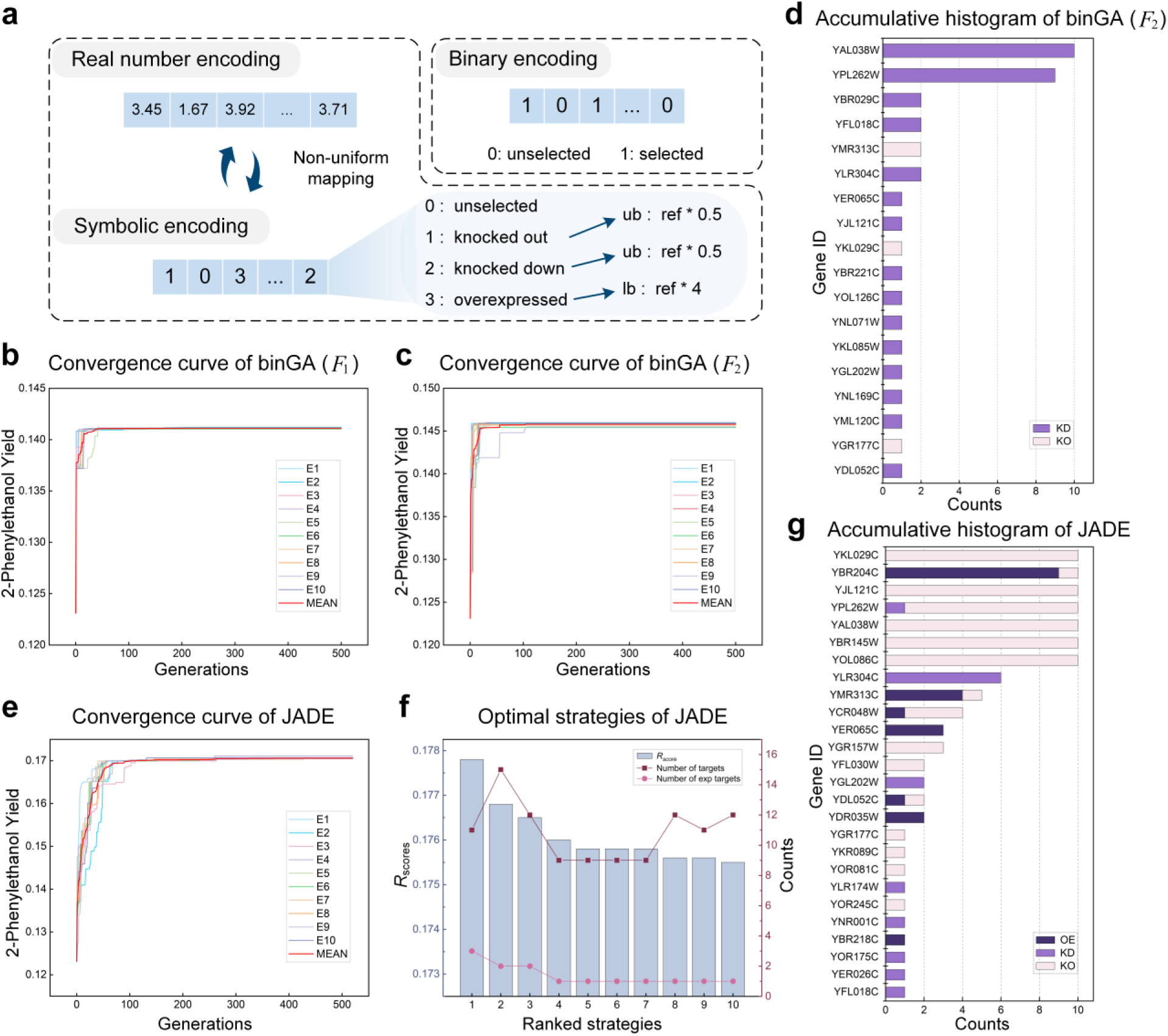
Identification of combinatorial targets for 2-phenylethanol (2-PE) production in *S. cerevisiae*. (a) MetaStrain has designed multiple individual encoding schemes, which are suitable for various optimization algorithms. (b, c) Convergence performance of the binGA search process. The plots compare the evolution of the population’s average fitness over generations (b) with and (c) without the inclusion of a regularization term penalizing the number of gene modifications. (d, g) Accumulative histograms summarizing the frequency of each gene’s appearance in the optimal solutions identified across ten independent runs. (e) Convergence performance of the JADE search process. (f) Detail information of optimal strategies predicted by JADE, including the comprehensive scores, number of targets and validated targets of each strategy.

**Table 1.** The result of ecFSEOF.

| Operation | 2-PE targets count | Spermidine targets count | L-tryptophan targets count |
| --- | --- | --- | --- |
| OE | 11 | 68 | 37 |
| KD | 35 | 0 | 20 |
| KO | 6 | 18 | 0 |
| Total | 52 | 86 | 57 |

Note: OE, overexpression; KD, knockdown; KO, knockout.

We first implemented binGA, which was performed on the above 52 candidate genes. The algorithm demonstrated robust convergence results for both fitness functions, with (*F*_2_) and without a regularization term (*F*_1_), identifying multiple biologically meaningful editing strategies (Fig. 2b, c). The binGA search using two fitness functions both employed elitism strategy. Among these, a strategy involving the knockdown of two targets—YAL038W (*CDC19*) and YPL262W (*FUM1*)—was repeatedly predicted (Fig. 2d). Mechanistically, *CDC19* encodes pyruvate kinase (Tripodi et al., 2015), a key enzyme in glycolysis that catalyzes the irreversible conversion of phosphoenolpyruvate to pyruvate. Knocking down *CDC19* reduces the glycolytic flux, thereby decreasing the input of pyruvate into the tricarboxylic acid (TCA) cycle, increasing the availability of the precursor phenylalanine and promoting 2-PE biosynthesis via the Ehrlich pathway (Dai et al., 2021). *FUM1* encodes fumarase (Wu and Tzagoloff, 1987), which catalyzes the conversion of fumarate to malate in the TCA cycle. Knocking down *FUM1* weakens TCA cycle activity, reducing excessive competition in energy metabolism and further optimizing carbon flux allocation toward the target product. The combined knockdown of *CDC19* and *FUM1* not only decreases resource consumption by competing metabolic pathways but also synergistically enhances carbon flux allocation to the target metabolic pathway. This strategy effectively reshapes global resource distribution in the metabolic network, providing a clear combinatorial target for efficient biosynthesis of 2-PE (Table 2). Additionally, non-intuitive edits such as knockdown *ARO8* and *ACO1* were also identified, offering testable hypotheses for experimental validation.

**Table 2.** Details of the top-ranked gene editing strategies predicted by binGA (*F_2_*)

| Algorithm | Individual<br>(Strategy) | Fitness<br>score | OE | KD | KO | Number<br>of targets | Number of<br>validated<br>targets |
| --- | --- | --- | --- | --- | --- | --- | --- |
| binGA<br>($F_2$) | $\mathbf{x}_1$ | 0.1470 | \ | **YAL038W**<br>YPL262W | \ | 2 | 1 |
| JADE | $\mathbf{x}_2$ | 0.1778 | **YDR035W**<br>YER065C<br>YBR204C | **YGL202W**<br>YLR304C | YOL086C<br>YBR145W<br>**YAL038W**<br>YPL262W<br>YJL121C<br>YKL029C | 11 | 3 |
| | $\mathbf{x}_3$ | 0.1768 | YBR218C<br>YDL052C<br>YCR048W | YBR145W<br>YFL018C<br>**YGL202W**<br>YLR304C<br>YFL030W<br>YOR245C | YOL086C<br>**YAL038W**<br>YPL262W<br>YJL121C<br>YBR204C<br>YKL029C | 15 | 2 |
| | $\mathbf{x}_4$ | 0.1765 | **YDR035W**<br>YER065C<br>YBR204C<br>YMR313C | YPL262W<br>YLR174W | YOL086C<br>YBR145W<br>**YAL038W**<br>YCR048W<br>YJL121C<br>YKL029C | 12 | 2 |

To unleash the full potential of *in silico* design, we removed the specified editing direction for each candidate gene and searched solely on the reduced gene set. This strategy significantly expanded the search space while increasing the complexity of the search landscape, imposing higher demands on algorithm performance. Initially, we employed symGA for the search. Despite incorporating an elite retention strategy, symGA demonstrated fast convergence (Fig. S3a, b). It exhibited low stability across ten independent experiments (Table S4), with no single target consistently predicted across all ten runs, and many genes exhibited diverse editing strategies in different combinations (Fig. S3c). This inconsistency shows the limitations of symGA in finding stable solution, but it can provide multiple satisfactory local optimal solutions. To overcome the bottleneck of symGA, we have chosen a high-performance algorithm called JADE. Across ten independent runs, JADE achieved stable convergence and produced highly consistent solutions (Fig. 2e, f). Statistical analysis revealed that seven genes recurred in all best-performing strategies with near-identical editing modes (Fig. 2g), which supports biological plausibility and facilitates experimental design. JADE’s adaptive parameter control further improved optimization efficiency by eliminating the need for manual parameter tuning. Details of the top-ranked strategies are reported in Table 2. Mechanistically, overexpressing *LDH1* (Chen et al., 2025) enhances the cytosolic reducing power needed for the NADH-dependent conversion of phenylacetaldehyde to 2-PE, while knocking out *CDC19* and *FUM1* throttles glycolytic outflow and TCA throughput, reallocating carbon and cofactors toward aromatic precursor formation. Deletion of *ADH1* and *ADH5* diminishes the ethanol sink (Thompson et al., 2018), preserving NADH and directing aldehyde reduction capacity to the Ehrlich branch. Additionally, deletion of *RPE1* and *MAE1* favoring erythrose-4-phosphate supply and redox balance for shikimate-pathway flux (Steyn et al., 2023). Together, these edits coherently rebalance carbon and redox to promote 2-PE synthesis.

### 3.3 Systematic performance analysis of meta-heuristic algorithms

We systematically evaluate the performance of the meta-heuristic algorithms within the MetaStrain framework on various products, focusing on convergence, stability, ability to control strategy size and cover experimented targets. Using *S. cerevisiae* and 2-PE as the primary testbed, we contrasted binGA that leverages annotation from the screening module with symGA and JADE operating in an expanded, annotation-free search space.

In the comparison of convergence performance across algorithms, the convergence curves of binGA, symGA (with and without an elite retention strategy), and JADE were plotted on the same graph (Fig. 3a), using *F*_1_ (Eq. (5)) as the fitness function. Results showed that binGA, with its simple search strategy, converged quickly. In contrast, JADE exhibited higher convergence quality due to its powerful global search capabilities, optimizing the objective function further in complex search landscapes. The convergence speed of symGA can be significantly accelerated by elite retention strategy, but the symGA showed poor stability in repeated experiments. Both binGA and JADE showed a relatively low standard deviation in the results of ten repeated experiments (Fig. 3b). Controlling the number of gene targets is critical for metabolic engineering, as excessive targets can increase experimental costs and reduce strain modification feasibility. We compared the target counts from optimal strategies generated independently by binGA (*F*_1_ and *F*_2_) and JADE (Fig. 3c). Results indicated that binGA combined with a regularization strategy effectively limited the number of predicted targets. In contrast, JADE, relying on its strong search performance, automatically excluded redundant targets during iterations, yielding strategies that were acceptable for wet-lab validation. Because symGA’s convergence is unstable, it was not included in the comparison. Notably, fewer targets do not necessarily lead to better strategies. Enforcing strict limits on target numbers may hinder the algorithm’s ability to explore potentially superior solutions, which could negatively impact rational MCFs design. MetaStrain provides effective mechanisms to balance solution complexity with performance, enhancing the feasibility and cost-efficiency of strain modifications.

**Fig. 3.**
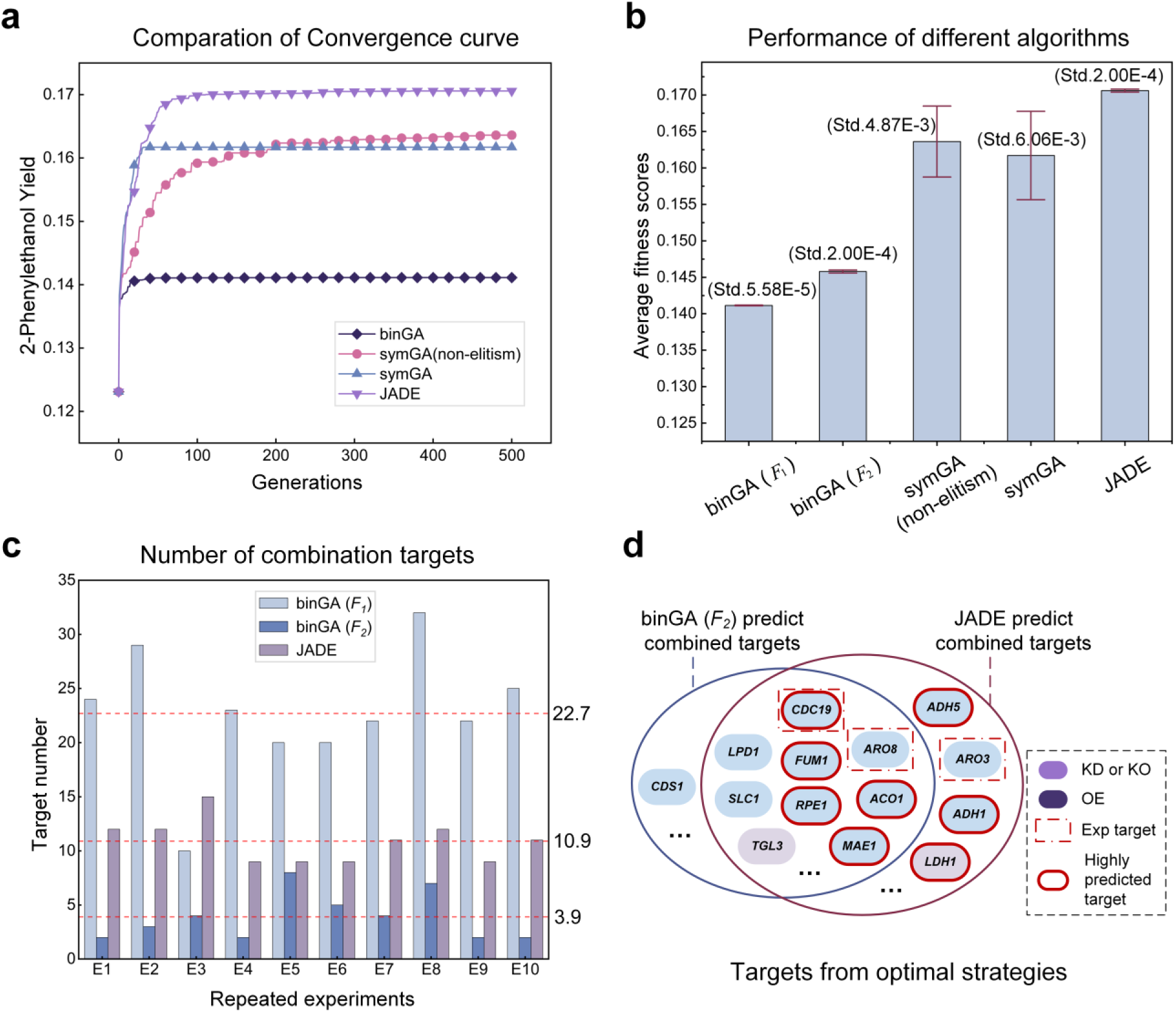
Comprehensive performance comparison and target analysis of different optimization algorithms. (a) Convergence profiles of the four algorithms evaluated using *F*_1_ in this study, illustrating their relative efficiency and effectiveness in reaching optimal solutions. (b) The convergence means and standard deviations of different algorithms. (c) A bar chart comparing the number of genetic modifications required by the optimal strategies across ten independent runs. This highlights the ability of some algorithms to find more experimentally feasible solutions. (d) A Venn diagram illustrating the overlap between the predicted gene targets and a set of experimentally validated targets from literature.

The effectiveness of metabolic engineering optimization algorithms hinges on their ability to identify experimentally validated gene targets while proposing innovative and insightful strategies. The results of binGA (*F*_2_) and JADE demonstrated that both algorithms frequently identified key targets. JADE further exhibited superior predictive performance, with its high-frequency targets encompassing those of binGA (Fig. 3d). Moreover, the overlapping gene editing strategies were consistent, such as the knockdown of *CDC19* (experimentally validated) and *FUM1*. Additionally, the screening performed by ecFSEOF ensured that the initial candidate pool included seven validated targets, with each strategy containing one to three validated targets, enhancing the biological reliability of the predictions. Beyond validated genes, both algorithms proposed numerous novel gene targets that have not been reported (Domenzain et al., 2025; Wei et al., 2022), exhibiting significant engineering value due to their potential roles in redistributing carbon flux or reducing network redundancy to enhance product yield. Such non-intuitive predictions highlight the power of meta-heuristic algorithms in optimizing complex networks, uncovering latent targets that conventional empirical methods fail to identify.

We further assessed generalization by switching the objective to spermidine overproduction in *S. cerevisiae*. Spermidine, a biologically significant polyamine compound with potential in anti-aging and cell growth regulation (Madeo et al., 2018), presents its own metabolic trade-offs (Fig. S2b). After ecFSEOF identified 86 potential gene targets (Table 1). We deployed JADE for this task, which again demonstrated stable convergence (Fig. S4a, b). Statistical analysis of ten independent runs revealed that two targets were consistently selected in all predictions with identical editing directions: the knockout of *EMI2* and the overexpression of *SPE3* (Fig. S4c, Table S5). *SPE3* (experimentally validated) encodes arginine decarboxylase, which catalyzes the conversion of arginine to ornithine—a crucial upstream step in spermidine biosynthesis (Friesen et al., 1998). Overexpression of *SPE3* significantly enhanced the supply of precursors for spermidine synthesis. Conversely, *EMI2* knockout mitigated competing ornithine catabolism, further improving carbon and nitrogen allocation. Additionally, JADE predicted other validated targets, such as *APT1* and *CAR2*, related to energy supply and nitrogen metabolism, respectively. The successful identification of both validated and novel synergistic targets (Table S6) in a completely different biosynthetic pathway underscores MetaStrain’s robustness and versatility as a general-purpose platform for accelerating the design of complex MCFs.

### 3.4 Metastrain prediction: single-target enhancement of L-tryptophan biosynthesis in *E. coli*

To evaluate the predictive performance of the MetaStrain algorithm, we focused on enhancing the production of L-tryptophan, an essential amino acid, by predicting single-target modifications. Predicted single targets were then validated using the initial L-tryptophan producing strain *E. coli* MGHT-10-1. First, the *E. coli* enzyme-constraint model eciML1515 (Ye et al., 2020) was refined to ensure that its genotype, culture medium environment, and baseline phenotype matched those of the experimental strain MGHT-10-1 (Materials and Methods). After model refinement, the simulated initial L-tryptophan production rate was 0.0111. Next, ecFSEOF was applied for dimensionality reduction, resulting in a set of 57 candidate genes (Table 1). Genes involved in the terminal steps of L-tryptophan biosynthesis were excluded to bias the algorithm towards identifying potential targets in other pathways that could synergistically regulate L-tryptophan production (Fig. 4a). In the search phase, the JADE optimization algorithm was employed with *F*_1_ as the fitness function for a global search of single-target modifications. To assess the stability of the algorithm and to generate diverse strategy recommendations, 20 independent replicate searches were performed (Fig. S5). The algorithm demonstrated good convergence and consistency, with a mean optimal fitness of 0.0449, including an average of 9 targets per search. Several genes were frequently predicted; for example, the *lpd* gene was identified as a knockdown target in all 20 searches. Each optimal strategy underwent further single-target redundancy analysis (Materials and Methods, Table S7), which confirmed that b0116, b4025, b3386, and b4094 were essential targets in multiple optimal strategies (Fig. 4b).

**Fig. 4.**
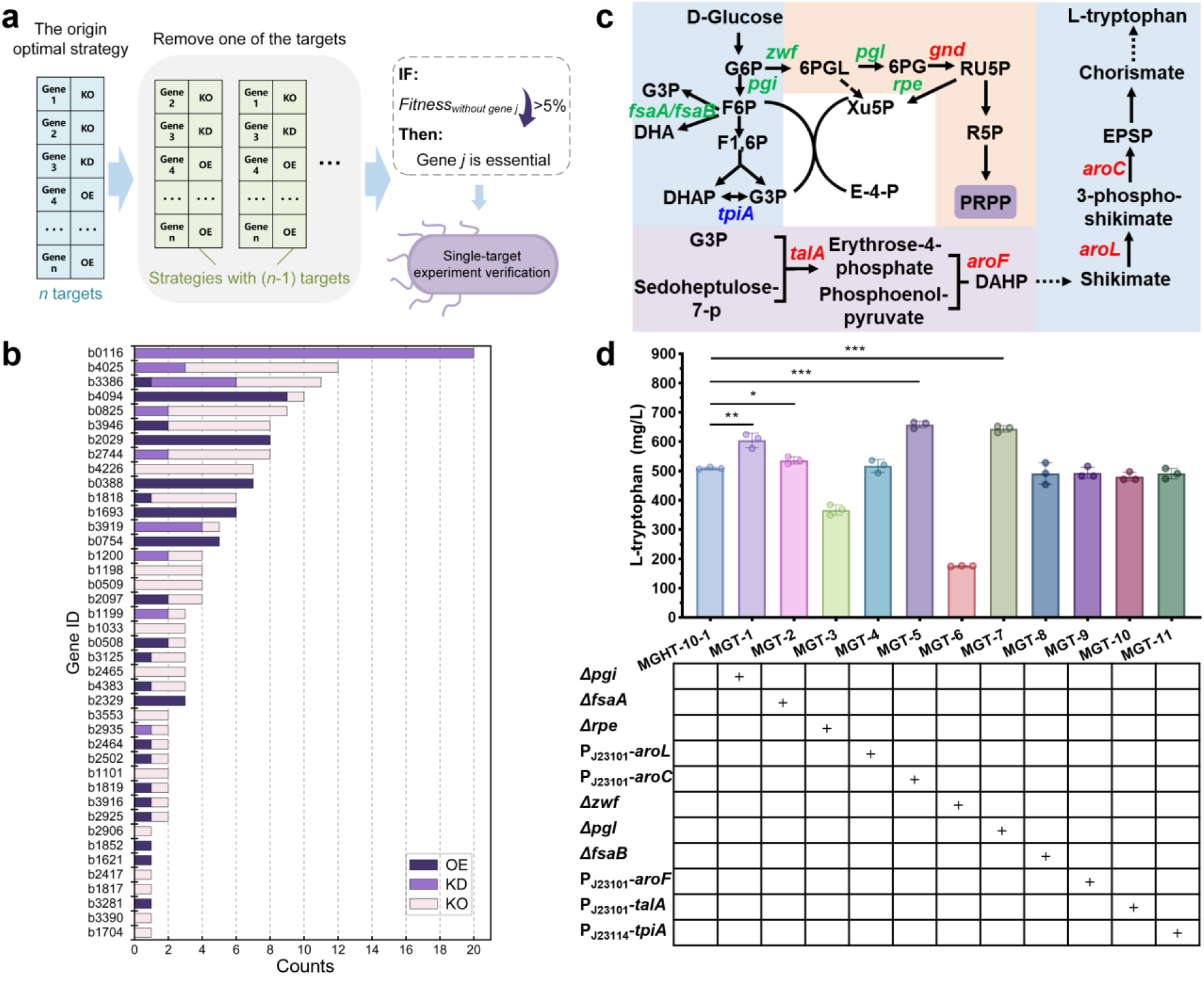
Single-target enhancement of L-tryptophan biosynthesis in *E. coli* by MetaStrain’s prediction. (a) Single-target prediction diagram by MetaStrain. (b) Counts of each gene in the 20 single-target prediction results. Overexpression (OE), Knockdown (KD), and Knockout (KO) denote these genetic perturbations. (c) L-tryptophan biosynthesis pathway schematic. (d) The impact of single targets predicted for engineering on L-tryptophan biosynthesis. *: *p* < 0.05, \*\**p*: < 0.01, and ***: *p* < 0.001. G6P: glucose-6-phosphate, F6P: fructose-6-phosphate, F1,6P: fructose-1,6-bisphosphate, G3P: glyceraldehyde 3-phosphate, DHAP: dihydroxyacetone phosphate, 6PG: 6-phosphogluconate, RU5P: ribulose-5-phosphate, R5P: ribose-5-phosphate, PRPP: phosphoribosyl pyrophosphate, XU5P: xylulose-5-phosphate, E-4-P: erythrose-4-phosphate, DHA: dihydroxyacetone, 6PGL: 6-Phosphogluconolactone.

Eleven of the predicted targets were selected for experimental validation. These targets are primarily distributed within the glycolysis pathway, the pentose phosphate pathway (PPP), and the shikimate pathway (Fig. 4c). The following genes were either knocked out, knocked down, or overexpressed in *E. coli* MGHT-10-1: *pgi* (b4025) encoding glucose-6-phosphate isomerase; *fsaA* (b0825) and *fsaB* (b3946) encoding fructose-6-phosphate aldolase; *tpiA* (b3919) encoding triosephosphate isomerase; *zwf* (b1852) encoding glucose-6-phosphate 1-dehydrogenase; *pgl* (b0767) encoding 6-phosphogluconolactonase; *gnd* (b2029) encoding 6-phosphogluconate dehydrogenase; *rpe* (b3386) encoding ribulose-phosphate 3-epimerase; *talA* (b2464) encoding transaldolase A; *aroF* (b2601) encoding 3-deoxy-7-phosphoheptulonate synthase; *aroL* (b0388) encoding shikimate kinase; and *aroC* (b2329) encoding chorismate synthase. Among the knockout targets, deletion of *pgi*, *fsaA*, and *pgl* significantly increased L-tryptophan production, with recombinant strains MGT-1, MGT-2, and MGT-7 reaching L-tryptophan titers of 604.57 mg/L, 536.07 mg/L, and 644.13 mg/L, respectively (Fig. 4d). These titers represent increases of 18.52%, 5.09%, and 26.28% compared to the control strain MGHT-10-1 (510.09 mg/L). Deletion of *fsaB* had little effect on L-tryptophan synthesis (strain MGT-8). However, deletion of *rpe* and *zwf* significantly reduced L-tryptophan production, with deletion of *zwf* (strain MGT-6) severely impacting *E. coli* growth (Fig. S6). For overexpression targets, the *aroL*, *aroC*, *aroF*, and *talA* genes were overexpressed using the strong constitutive promoter P_J23101_, resulting in recombinant strains MGT-4, MGT-5, MGT-9, and MGT-10. Overexpression of *aroC* promoted L-tryptophan synthesis, achieving a titer of 658.52 mg/L. The L-tryptophan titers of the other overexpression strains were not significantly different from that of the control strain MGHT-10-1. For knockdown targets, the expression of *tpiA* was weakened using the weak promoter P_J23114_, but this had little effect on L-tryptophan production. In summary, of the 11 single targets predicted by the MetaStrain algorithm and tested experimentally, 4 significantly increased the production of L-tryptophan.

### 3.5 Combinatorial target engineering optimizes L-tryptophan yield in *E. coli*

A core strength of the MetaStrain framework is its ability to account for the non-linear combinatorial effects among gene targets. To experimentally validate these predicted synergies, we adopted a screening strategy with a fixed number of four targets per combination for strain construction (Materials and Methods, Fig. 5a). The *in silico* performance evaluations for four- and five-target combinations are provided in the Table S8 and Table S9, with the top eight combinations shown in Fig. 5b. From these, four four-target combinations were selected for wet-lab validation. The results revealed that engineered strains carrying combinations that included a *pgi* knockout exhibited markedly increased biomass (Fig. 5c), including intermediate two-, three-, and four-target recombinant strains (MGT-12, MGT-14, and MGT-16).

**Fig. 5.**
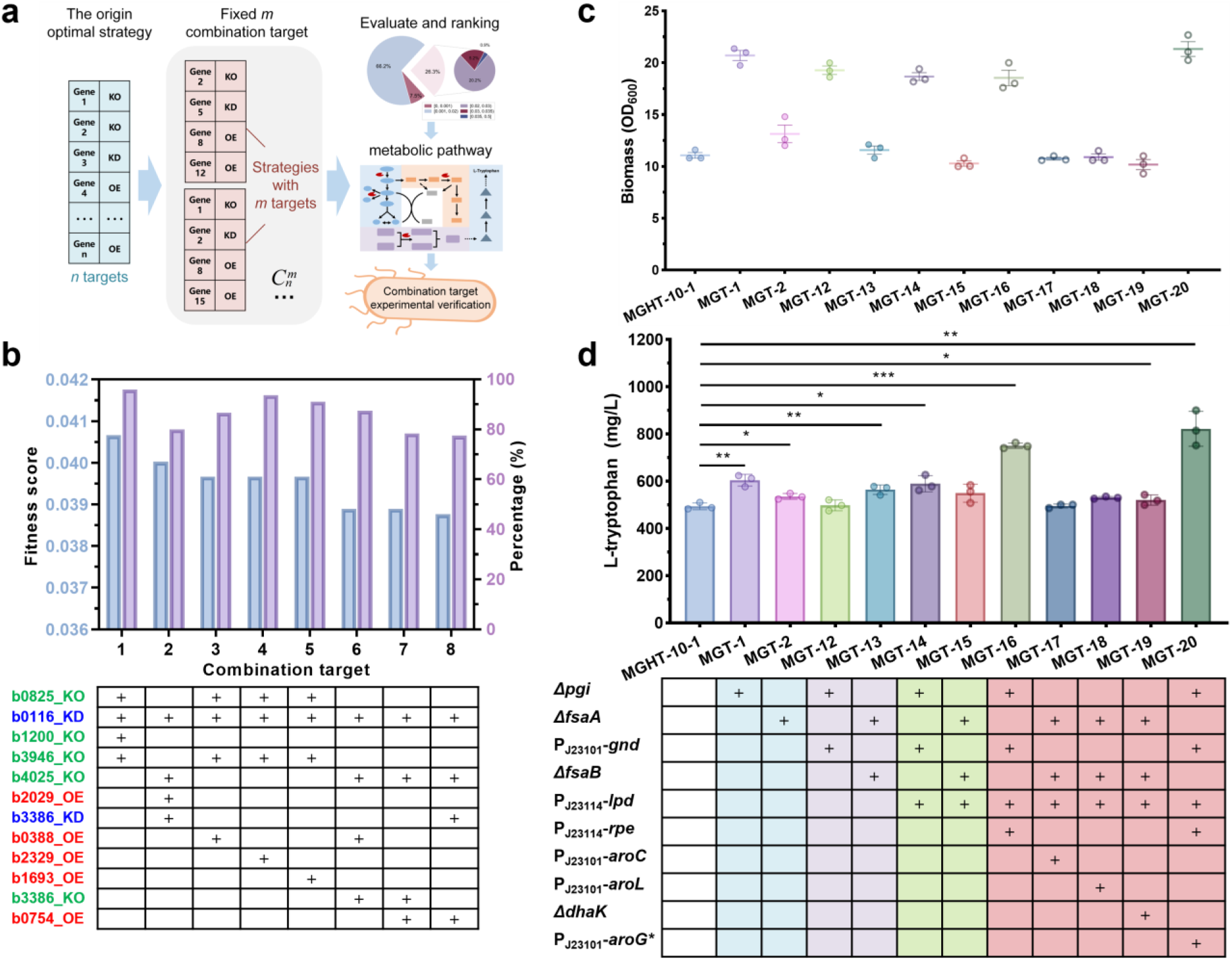
MetaStrain-predicted combinatorial targets for optimizing L-tryptophan production in *E. coli*. (a) Combinatorial targets prediction diagram by MetaStrain. (b) Top eight predicted four-target combinations. (c) Biomass of combinatorial target strains. (d) L-tryptophan production of combinatorial target strains. *: *p* < 0.05, **: *p* < 0.01, and ***: *p* < 0.001.

To investigate the stepwise combinatorial effects, intermediate strains with two or three targets were constructed and tested. Among the two-target combinations, strain MGT-13 (*fsaA*Δ *fsaB*Δ) achieved a titer of 564.84 mg/L, representing a 10.7% increase over the control. In contrast, strain MGT-12 (*pgi*Δ- P_J23101_-*gnd*) showed no improvement in L-tryptophan production. For the three-target combinations, strain MGT-14 (*pgi-*ΔP_J23101_-*gnd-*P_J23114_-*lpd*) yielded 589.52 mg/L of L-tryptophan, a 15.57% increase compared to the control. However, the L-tryptophan titer of strain MGT-15 (*fsaA*Δ-*fsaB*Δ-P_J23114_-*lpd*) did not improve. Among the four validated four-target combinations, two strains demonstrated a significant increase in L-tryptophan titer compared to the control strain MGHT-10-1. Notably, the engineered strain MGT-16 (*pgi*Δ-P_J23101_-*gnd*- P_J23114_-*lpd*-P_J23114_-*rpe*) achieved a titer of 751.02 mg/L, a remarkable 47.23% improvement over the control. Strain MGT-18 also showed a modest increase to 531.06 mg/L (a 4.11% improvement) (Fig. 5d). However, the other two four-target recombinant strains, MGT-17 and MGT-19, did not show any improvement in L-tryptophan production. In summary, of the four distinct four-target combinations predicted by MetaStrain’s strategy, two resulted in an increased L-tryptophan yield (Fig. 5d). Similarly, two of the four tested intermediate (two- and three-target) strains also showed significantly higher titers than the control strain. Subsequently, the effect of the five-target combination was further verified. Based on the strain MGT-16, the expression of *aroG\** (S180F) was overexpressed to obtain the recombinant strain MGT-20. The results showed that the yield of L-tryptophan was 822.53 mg/L. It increased by 61.25% compared to the control strain MGT-10-1. These results powerfully demonstrate the efficacy of our combinatorial design approach. The 61.25% yield enhancement achieved by the MGT-20 strain far surpasses the improvements from any of the single-gene modifications, providing definitive experimental evidence for the synergistic effects captured by the MetaStrain algorithm. The successful engineering of superior producers through these non-intuitive combinations—which involve coordinated adjustments across glycolysis, the PPP, and cofactor-related pathways—highlights the framework’s ability to navigate vast and complex design spaces. This work not only validates MetaStrain as a powerful tool for accelerating the metabolic engineering cycle but also underscores the immense potential of integrating advanced search algorithms with genome-scale models to unlock complex phenotypes that lie beyond the reach of rational design and single-target intuition.

## 4. Discussion

The MetaStrain algorithm proposed in this study combines ecModels with efficient meta-heuristic algorithms, offering an innovative solution to the combinatorial target optimization challenges in complex metabolic engineering. Its exceptional performance across multiple optimization tasks demonstrates the algorithm’s ability to identify non-intuitive and innovative gene editing strategies while ensuring result stability, thus providing critical support for developing efficient microbial cell factories. While GA provide simplicity and efficiency, particularly in smaller search spaces, the JADE significantly enhances global search capability through adaptive parameter control and external archive mechanisms, achieving efficient convergence even in large and complex search spaces. Notably, JADE accurately identifies experimentally validated targets and proposes novel targets, underscoring MetaStrain’s potential in integrating existing knowledge with uncovering new strategies. The strength of MetaStrain lies in its ability to transition from single-target to combinatorial-target optimization, providing a comprehensive means of addressing complex metabolic network competition and reshaping carbon flux distribution. The MetaStrain algorithm has been successfully applied to predict the target genes for the synthesis of 2-PE and spermidine in *S. cerevisiae*. Moreover, it has also been used to predict the single-targets and combined-targets of L-tryptophan in *E. coli* and significantly enhance its production. Its successful adaptation to the replacement of the target product further reinforces its broad applicability.

In the prediction of *S. cerevisiae* for the synthesis of 2-PE and spermidine, the MetaStrain algorithm identified combination targets that included three experimentally validated genes (*CDC19*, *ARO8*, and *ARO3*) and one validated gene (*SPE3*), respectively. For the prediction of single-gene targets in *E. coli* for L-tryptophan production, experimental validation confirmed 11 target genes, four of which significantly enhanced L-tryptophan yield. Among these, the knockout of *pgi* was particularly notable, as it substantially increased both biomass and L-tryptophan production in *E. coli*. The observed improvement in biomass following *pgi* deletion is consistent with prior findings (Liu et al., 2025a). The increase in L-tryptophan yield is likely due to the redirection of carbon flux from glycolysis into the pentose phosphate pathway (PPP) upon *pgi* disruption, thereby augmenting the supply of precursors such as ribose-5-phosphate, erythrose-4-phosphate, and the cofactor NADPH.

In the case of combinatorial target predictions for L-tryptophan production in *E. coli*, the most effective combination was *pgi*Δ-P_J23101_-*gnd*-P_J23114_-*lpd*-P_J23114_-*rpe-*P_J23101_-*aroG\**, which resulted in a 61.25% increase in yield compared to the control strain. Within this combination, attenuation of *lpd* is proposed to reduce the flux of pyruvate toward acetyl-CoA (Aspacio et al., 2024; Bouzon et al., 2021), potentially leading to accumulation of the L-tryptophan precursor phosphoenolpyruvate and thus promoting synthesis. Overall, among the predicted combinatorial targets, approximately 50% were confirmed to enhance L-tryptophan synthesis. In conclusion, MetaStrain algorithm provides reasonably accurate gene target predictions for the construction of microbial cell factories. Despite MetaStrain’s promising performance, there remains room for improvement.

Incorporating multi-objective optimization algorithms could enable simultaneous optimization of multiple indicators, enhancing practical applicability. Additionally, integrating surrogate models, such as machine learning-based metabolic flux prediction models, could accelerate the computational process, particularly in large-scale search spaces. Exploring the integration of more meta-heuristic algorithms and dynamic environment optimization mechanisms may further enhance the algorithm’s adaptability and robustness.

In conclusion, the MetaStrain algorithm offers a novel chance for optimizing complex metabolic networks, with its innovation, stability, and adaptability, laying a solid foundation for advancing metabolic engineering. Combining MetaStrain with high-throughput experimental platforms could extend its impact from computational optimization to experimental validation, creating a closed-loop design process that tightly couples computation and experimentation. This synergistic approach will not only accelerate the development of microbial cell factories but also drive the widespread application of metabolic engineering in sustainable biomanufacturing solutions.

## Supporting information

Supplementary Material for MetaStrain

## CRediT authorship contribution statement

Wenbin Liao: Writing – review & editing, Writing – original draft, Visualization, Investigation, Designing and debugging algorithms. Gengrong Gao: Writing – review & editing, Writing – original draft, Visualization, Investigation, Experimental verification. Haoyu Wang: Investigation. Xingcun Fan: Investigation. Siwei He: Investigation. Guangming Xiang: Investigation. Xuefeng Yan: Writing – review & editing, Supervision, Funding acquisition. Hongzhong Lu: Writing – review & editing, Supervision, Funding acquisition.

## Acknowledgement

This work was financially supported by the National key research and development program of China (2020YFA0908300), Shanghai Municipal Science and Technology Major Project, and grant 22208211 and 22378263 from the National Natural Science Foundation of China (NSFC).

## Data and code availability

The MetaStrain method is available as a standalone tool at the public repository: https://github.com/MBLiao/MetaStrain.

## Declaration of generative AI and AI-assisted technologies in the manuscript preparation process

During the preparation of this work the authors used Chat-GPT 4 in order to refine the language of the article. After using this tool, the authors reviewed and edited the content as needed and take full responsibility for the content of the published article.

## References

Aspacio, D., Luu, E., Worakaensai, S., Cui, Y., Maxel, S., King, E., Hagerty, R., Chu, A., Minn, D., Siegel, J. B., Li, H., 2024. Engineering *Escherichia coli* Pyruvate Metabolism to Generate Noncanonical Reducing Power. ACS Catal. 14, 9776–9784.

Basan, M., Hui, S., Okano, H., Zhang, Z., Shen, Y., Williamson, J. R., Hwa, T., 2015. Overflow metabolism in *Escherichia coli* results from efficient proteome allocation. Nature. 528, 99–104.

Beg, Q. K., Vazquez, A., Ernst, J., de Menezes, M. A., Bar-Joseph, Z., Barabasi, A. L., Oltvai, Z. N., 2007. Intracellular crowding defines the mode and sequence of substrate uptake by *Escherichia coli* and constrains its metabolic activity. Proc Natl Acad Sci U S A. 104, 12663–8.

Bouzon, M., Döring, V., Dubois, I., Berger, A., Stoffel, G. M. M., Calzadiaz Ramirez, L., Meyer, S. N., Fouré, M., Roche, D., Perret, A., Erb, T. J., Bar-Even, A., Lindner, S. N., 2021. Change in Cofactor Specificity of Oxidoreductases by Adaptive Evolution of an *Escherichia coli* NADPH-Auxotrophic Strain. mBio. 12, e0032921.

Burgard, A., Pharkya, P., Maranas, C., 2003. OptKnock: A bilevel programming framework for identifying gene knockout strategies for microbial strain optimization. Biotechnology and Bioengineering. 84, 647–657.

Chen, Y., Tang, J., Li, Q., Xin, W., Xiao, X., Mou, B., Li, J., Lu, F., Fu, C., Long, W., Liao, H., Han, X., Feng, P., Li, W., Zhou, K., Yang, L., Chen, X., Yang, L., Ma, M., Yang, Y., Wang, H., 2025. Analysis of the response mechanism in lipid degradation of key gene LDH1 knockout strains of *Saccharomyces cerevisiae* under levulinic acid stress. Arch Microbiol. 207, 188.

Choi, H. S., Lee, S. Y., Kim, T. Y., Woo, H. M., 2010. *In silico* identification of gene amplification targets for improvement of lycopene production. Appl Environ Microbiol. 76, 3097–105.

Dai, J., Xia, H., Yang, C., Chen, X., 2021. Sensing, Uptake and Catabolism of L-Phenylalanine During 2-Phenylethanol Biosynthesis via the Ehrlich Pathway in *Saccharomyces cerevisiae*. Front Microbiol. 12, 601963.

Das, S., Suganthan, P. N., 2011. Differential Evolution: A Survey of the State-of-the-Art. Ieee Transactions on Evolutionary Computation. 15, 4–31.

Deb, K., Pratap, A., Agarwal, S., Meyarivan, T., 2002. A fast and elitist multiobjective genetic algorithm: NSGA-II. IEEE Transactions on Evolutionary Computation. 6, 182–197.

Domenzain, I., Lu, Y., Wang, H., Shi, J., Lu, H., Nielsen, J., 2025. Computational biology predicts metabolic engineering targets for increased production of 103 valuable chemicals in yeast. Proc Natl Acad Sci U S A. 122, e2417322122.

Duman-Özdamar, Z. E., Binay, B., 2021. Production of Industrial Enzymes via Pichia pastoris as a Cell Factory in Bioreactor: Current Status and Future Aspects. Protein J. 40, 367–376.

Friesen, H., Tanny, J. C., Segall, J., 1998. Spe3, which encodes spermidine synthase, is required for full repression through NRE(DIT) in *Saccharomyces cerevisiae*. Genetics. 150, 59–73.

Gurobi Optimization, L. L. C., Gurobi Optimizer Reference Manual.

Holland, J. H., 1992. Genetic Algorithms. Scientific American. 267, 66-72.

Huang, C., Li, Y., Yao, X., 2020. A Survey of Automatic Parameter Tuning Methods for Metaheuristics. IEEE Transactions on Evolutionary Computation. 24, 201–216.

Jensen, K., Broeken, V., Hansen, A. S. L., Sonnenschein, N., Herrgard, M. J., 2019. OptCouple: Joint simulation of gene knockouts, insertions and medium modifications for prediction of growth-coupled strain designs. Metabolic engineering communications. 8, e00087–e00087.

Jiang, S., Otero-Muras, I., Banga, J. R., Wang, Y., Kaiser, M., Krasnogor, N., 2022. OptDesign: Identifying Optimum Design Strategies in Strain Engineering for Biochemical Production. Acs Synthetic Biology. 11, 1531–1541.

Jullesson, D., David, F., Pfleger, B., Nielsen, J., 2015. Impact of synthetic biology and metabolic engineering on industrial production of fine chemicals. Biotechnology Advances. 33, 1395–1402.

Konzock, O., Nielsen, J., 2024. TRYing to evaluate production costs in microbial biotechnology. Trends Biotechnol. 42, 1339–1347.

Lewis, N. E., Nagarajan, H., Palsson, B. O., 2012. Constraining the metabolic genotype-phenotype relationship using a phylogeny of *in silico* methods. Nature Reviews Microbiology. 10, 291–305.

Li, Q., Sun, B., Chen, J., Zhang, Y., Jiang, Y., Yang, S., 2021. A modified pCas/pTargetF system for CRISPR-Cas9-assisted genome editing in *Escherichia coli*. Acta Biochim Biophys Sin (Shanghai). 53, 620–627.

Liu, L., Zhang, X., Zhu, T., Ye, T., Ding, W., Liu, H., Fang, H., 2025a. Combined Transcriptomics and (13)C Metabolomics Analysis Reveals pgi and edd Genes Involved in the Regulation of Efficient Cytidine Synthesis in *Escherichia coli*. ACS Synth Biol. 14, 542–552.

Liu, W., Zhang, Y., Liu, K., Quinn, B., Yang, X., Peng, Q., 2025b. Evolutionary Multiobjective Optimization for Large-Scale Portfolio Selection With Both Random and Uncertain Returns. IEEE Transactions on Evolutionary Computation. 29, 76–90.

Lu, H., Li, F., Sanchez, B. J., Zhu, Z., Li, G., Domenzain, I., Marcisauskas, S., Anton, P. M., Lappa, D., Lieven, C., Beber, M. E., Sonnenschein, N., Kerkhoven, E. J., Nielsen, J., 2019. A consensus *S. cerevisiae* metabolic model Yeast8 and its ecosystem for comprehensively probing cellular metabolism. Nat Commun. 10, 3586.

Lu, J., Bi, X., Liu, Y., Lv, X., Li, J., Du, G., Liu, L., 2023. *In silico* cell factory design driven by comprehensive genome-scale metabolic models: development and challenges. Systems Microbiology and Biomanufacturing. 3, 207–222.

Madeo, F., Eisenberg, T., Pietrocola, F., Kroemer, G., 2018. Spermidine in health and disease. Science. 359.

Monk, J. M., Lloyd, C. J., Brunk, E., Mih, N., Sastry, A., King, Z., Takeuchi, R., Nomura, W., Zhang, Z., Mori, H., Feist, A. M., Palsson, B. O., 2017. iML1515, a knowledgebase that computes *Escherichia coli* traits. Nat Biotechnol. 35, 904–908.

Nielsen, J., Keasling, J. D., 2016. Engineering Cellular Metabolism. Cell. 164, 1185–1197.

Nielsen, J., Larsson, C., van Maris, A., Pronk, J., 2013. Metabolic engineering of yeast for production of fuels and chemicals. Curr Opin Biotechnol. 24, 398–404.

Nilsson, A., Nielsen, J., 2016. Metabolic Trade-offs in Yeast are Caused by F1F0-ATP synthase. Scientific Reports. 6.

O’Brien, E. J., Monk, J. M., Palsson, B. O., 2015. Using Genome-scale Models to Predict Biological Capabilities. Cell. 161, 971–987.

Orth, J. D., Thiele, I., Palsson, B. O., 2010. What is flux balance analysis? Nature Biotechnology. 28, 245–248.

Patil, K. R., Rocha, I., Forster, J., Nielsen, J., 2005. Evolutionary programming as a platform for *in silico* metabolic engineering. BMC Bioinformatics. 6, 308.

Ranganathan, S., Suthers, P. F., Maranas, C. D., 2010. OptForce: an optimization procedure for identifying all genetic manipulations leading to targeted overproductions. PLoS Comput Biol. 6, e1000744.

Ro, D. K., Paradise, E. M., Ouellet, M., Fisher, K. J., Newman, K. L., Ndungu, J. M., Ho, K. A., Eachus, R. A., Ham, T. S., Kirby, J., Chang, M. C., Withers, S. T., Shiba, Y., Sarpong, R., Keasling, J. D., 2006. Production of the antimalarial drug precursor artemisinic acid in engineered yeast. Nature. 440, 940–3.

Sabzevari, M., Szedmak, S., Penttila, M., Jouhten, P., Rousu, J., 2022. Strain design optimization using reinforcement learning. Plos Computational Biology. 18.

Sanchez, B. J., Zhang, C., Nilsson, A., Lahtvee, P.-J., Kerkhoven, E. J., Nielsen, J., 2017. Improving the phenotype predictions of a yeast genome-scale metabolic model by incorporating enzymatic constraints. Molecular Systems Biology. 13.

Schneider, P., Bekiaris, P. S., von Kamp, A., Klamt, S., 2022. StrainDesign: a comprehensive Python package for computational design of metabolic networks. Bioinformatics. 38, 4981–4983.

Segrè, D., Vitkup, D., Church, G., 2002. Analysis of optimality in natural and perturbed metabolic networks. Proceedings of the National Academy of Sciences of the United States of America. 99, 15112–15117.

Shen, T., Liu, Q., Xie, X., Xu, Q., Chen, N., 2012. Improved production of tryptophan in genetically engineered *Escherichia coli* with TktA and PpsA overexpression. J Biomed Biotechnol. 2012, 605219.

Shlomi, T., Benyamini, T., Gottlieb, E., Sharan, R., Ruppin, E., 2011. Genome-Scale Metabolic Modeling Elucidates the Role of Proliferative Adaptation in Causing the Warburg Effect. Plos Computational Biology. 7.

Steyn, A., Viljoen-Bloom, M., Van Zyl, W. H., 2023. Constructing recombinant *Saccharomyces cerevisiae* strains for malic-to-fumaric acid conversion. FEMS Microbiol Lett. 370.

Thompson, C. E., Freitas, L. B., Salzano, F. M., 2018. Molecular evolution and functional divergence of alcohol dehydrogenases in animals, fungi and plants. Genet Mol Biol. 41, 341–354.

Tripodi, F., Nicastro, R., Reghellin, V., Coccetti, P., 2015. Post-translational modifications on yeast carbon metabolism: Regulatory mechanisms beyond transcriptional control. Biochim Biophys Acta. 1850, 620–7.

Volk, M. J., Tran, V. G., Tan, S.-., I, Mishra, S., Fatma, Z., Boob, A., Li, H., Xue, P., Martin, T. A., Zhao, H., 2023. Metabolic Engineering: Methodologies and Applications. Chemical Reviews. 123, 5521–5570.

Wang, H., Zhang, M., Zhang, C., He, S., Liao, W., Zhu, R., Zhou, Y. J., Lu, H., 2025. StrainOptimizer empowers rational cell factory design through multi-scale metabolic models with expression and proteome constraints. bioRxiv.

Wang, X., Wang, Y., Tang, L., Zhang, Q., 2024. Multiobjective Ensemble Learning With Multiscale Data for Product Quality Prediction in Iron and Steel Industry. IEEE Transactions on Evolutionary Computation. 28, 1099–1113.

Wang, Y., Zhang, H., Lu, X., Zong, H., Zhuge, B., 2019. Advances in 2-phenylethanol production from engineered microorganisms. Biotechnol Adv. 37, 403–409.

Wei, Y., Bergenholm, D., Gossing, M., Siewers, V., Nielsen, J., 2018. Expression of cocoa genes in *Saccharomyces cerevisiae* improves cocoa butter production. Microb Cell Fact. 17, 11.

Wei, Y., Hao, G., Yujia, J., Xiujuan, Q., Jie, Z., Weiliang, D., Wenming, Z., Fengxue, X., Min, J., 2022. Research progress in 2-phenylethanol production through biological processes. Synthetic Biology Journal. 2, 1030–1045.

Wu, M., Tzagoloff, A., 1987. Mitochondrial and cytoplasmic fumarases in *Saccharomyces cerevisiae* are encoded by a single nuclear gene FUM1. J Biol Chem. 262, 12275–82.

Xiong, B., Zhu, Y., Tian, D., Jiang, S., Fan, X., Ma, Q., Wu, H., Xie, X., 2021. Flux redistribution of central carbon metabolism for efficient production of l-tryptophan in *Escherichia coli*. Biotechnol Bioeng. 118, 1393–1404.

Ye, C., Luo, Q., Guo, L., Gao, C., Xu, N., Zhang, L., Liu, L., Chen, X., 2020. Improving lysine production through construction of an *Escherichia coli* enzyme-constrained model. Biotechnol Bioeng. 117, 3533–3544.

Yu, Y., Tang, Q., Jiang, Q., Fan, Q., 2025. A Deep Reinforcement Learning-Assisted Multimodal Multi-Objective Bi-Level Optimization Method for Multi-Robot Task Allocation. IEEE Transactions on Evolutionary Computation. 1–1.

Zhang, J., Petersen, S. D., Radivojevic, T., Ramirez, A., Perez-Manriquez, A., Abeliuk, E., Sanchez, B. J., Costello, Z., Chen, Y., Fero, M. J., Martin, H. G., Nielsen, J., Keasling, J. D., Jensen, M. K., 2020. Combining mechanistic and machine learning models for predictive engineering and optimization of tryptophan metabolism. Nature Communications. 11.

Zhang, J., Sanderson, A. C., 2009. JADE: Adaptive Differential Evolution With Optional External Archive. IEEE Transactions on Evolutionary Computation. 13, 945–958.

Zhou, Y. J., Buijs, N. A., Zhu, Z., Qin, J., Siewers, V., Nielsen, J., 2016. Production of fatty acid-derived oleochemicals and biofuels by synthetic yeast cell factories. Nat Commun. 7, 11709.

