## Supplementary Material for MetaStrain for "Efficient Cell Factory Design by Combining Meta-Heuristic Algorithm with Enzyme-Constrained Metabolic Models"

### The ecFSEOF method

Initially, the modified algorithm sets the biomass synthesis flux to its theoretical maximum and progressively decreases it across 16 levels, ranging from 100% to 25% of the predicted maximum (Domenzain et al., 2025; Wang et al., 2025). This allows the algorithm to explore metabolic reaction changes under varying biomass synthesis conditions. FBA is then performed with the target product yield set as the objective function to obtain the flux values for all reactions. Flux changes are quantified through a two-step scoring process to identify reactions with consistent increases or decreases across these conditions. These reactions are considered potential engineering targets.

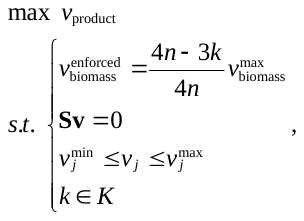

Here,
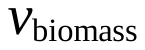
 represents the biomass synthesis flux in the ecModel. The algorithm incrementally reduces
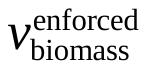
 in
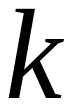
 steps, where
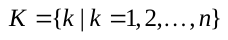
 (in this study
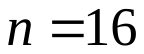
).

For each reaction, the flux score is calculated using the following formula:

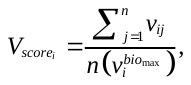

Where
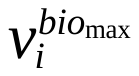
 is the flux of reaction under the maximum biomass synthesis rate condition, and
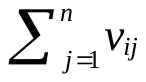
 is the sum of fluxes for reaction
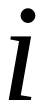
 across all simulation conditions, and
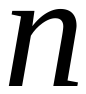
 is the number of simulations. The flux score is recalculated to account for potential metabolic mode shifts, such as transitions from respiratory to mixed fermentation metabolism. Flux values exceeding 1000 are capped at 1000, and undefined values (e.g., 0/0) are set to 1. Next, the gene score (K-score) is calculated as follows:

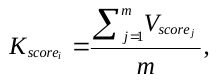

Here,
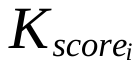
 is the score of gene target
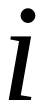
,
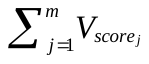
 is the score for each reaction catalyzed by the gene product, and
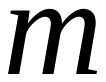
 is the total number of reactions catalyzed by the gene product. Based on the
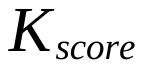
:

a) Genes with scores greater than 1 are suggested as overexpression targets.
b) Genes with scores between 0.05 and 0.5 are recommended as knockdown targets.
c) Genes with scores below 0.05 are proposed as knockout targets.

### Initialization and coding

The algorithm begins by generating a random initial population, which represents a set of potential solutions. Each individual corresponds to a point in the solution space. During the initialization phase, the population is generated as follows:

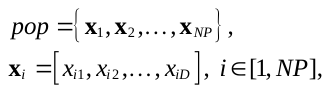

Here,
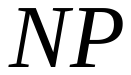
 represents the population size and
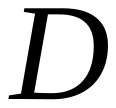
 indicates the dimension of each individual, corresponding to the number of genes selected after dimension reduction using ecFSEOF.

For binGA, each individual’s gene representation is binary, where “0” indicates not selected and “1” indicates selected. The initialization for binGA is:

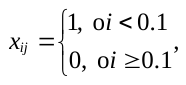

Here,
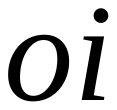
 is a random number between 0 and 1. The threshold of 0.1 ensures that fewer "1"s appear in the sequence.

For symGA, each gene has four possible states: unselected (“0”), knocked out (“1”), knocked down (“2”), and overexpressed (“3”). The initialization process is as follows:

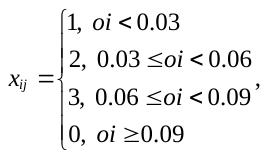

Genes are initialized with equal probability for one of the three editing states, but a higher probability is assigned for the gene to remain unselected.

For JADE, as the differential evolution process operates in a continuous decision space, each dimension of the individual is initialized by
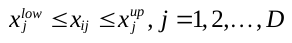
 with a random value within
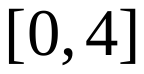
. Before evaluation, individuals are converted to discrete encodings:

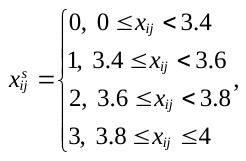

Here,
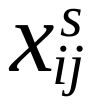
 represents the discretized individual, and non-uniform mapping effectively controls the number of editing targets in the strategy.

These initialization strategies ensure that the gene editing operations are feasible and avoid excessive interventions, making the solution economically viable and experimentally feasible.

### GA operators

The core principle of GA(Holland, 1992) is “survival of the fittest”. Through crossover, mutation, and selection, GA iteratively searches a population to find satisfactory solutions. The algorithm’s basic structure is illustrated in the corresponding flowchart.

In the selection stage, a roulette wheel selection strategy is employed. After evaluating each individual’s fitness, the selection probability for each individual is calculated as:

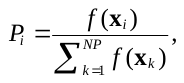

Here,
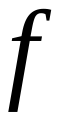
 represents the fitness value of individual
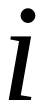
, obtained via ecMOMA. This formula computes the cumulative selection probability for all individuals. A random number between 0 and 1 is generated, and the individual corresponding to the interval in which this random number falls is selected. These selected individuals serve as candidates for crossover operations, with the number of participants adjustable based on the problem scale.

Crossover and mutation operations are both conducted probabilistically. In this study, single-point crossover is used. For binGA, mutation involves randomly selecting a gene locus in the individual and flipping its value (“0” to “1” or “1” to “0”).

For symGA, mutation is performed as follows:

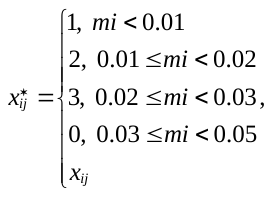

Here,
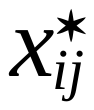
 represents the mutated value at the gene locus. The mutation direction threshold design ensures that genes undergoing mutation are more likely to remain unedited, effectively reducing the number of combinatorial targets in the results.

An elite retention strategy preserves the top 2–5% of individuals, bypassing crossover and mutation for direct propagation to the next generation. This accelerates convergence. After each iteration, the original population is mixed with newly generated individuals. The top individuals, based on fitness, are selected to form the next generation, ensuring continuous improvement.

### JADE operators

DE represent solutions using floating-point encoded individuals. Their process resembles that of GAs, but JADE(Zhang and Sanderson, 2009) introduces the following innovations in its optimization strategy:

a) Adaptive Parameter Control;
b) Optional External Archive;
c) Enhanced Mutation Strategy

After initialization, DE iterates through mutation, crossover, and selection until convergence. The external archive is initialized as empty. During each iteration, individuals eliminated in the selection process are added to the archive. If the archive size exceeds a predefined threshold (typically equal to the population size), some individuals are randomly removed to maintain its size. The archive provides additional information for the mutation process, enhancing search directions and further increasing diversity.

For generation
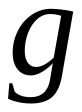
, the mutation operation generates a vector
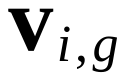
 for each parent individual. JADE uses a novel mutation strategy (
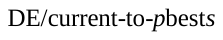
) defined as:

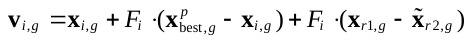

Here,
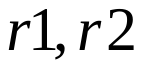
 are independent random variables and
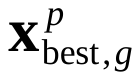
 is randomly selected from the top
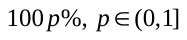
 of the population, while
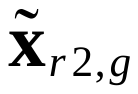
 randomly chosen from the union of the archive and current population. This
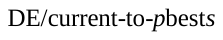
 strategy increases diversity without restricting the mutation process to a single optimal solution.

New individuals may exceed problem boundary constraints. A common solution is boundary control, where out-of-bound dimensions are reset to:

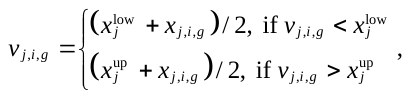

Here,
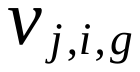
 is the
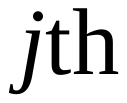
 dimensional variable of individual
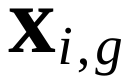
 of generation
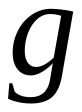
.This approach works well when the optimal solution lies near or at the boundary.

Crossover follows the mutation process, producing offspring
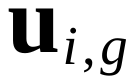

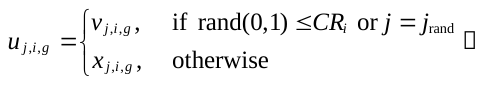

Here,
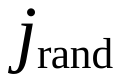
 is a random number between 0 and
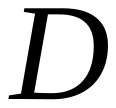
. In JADE, the crossover rate
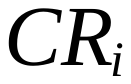
 was associated with each individual and was continuously updated during the search.

Selection retains the better individual between parent and offspring:

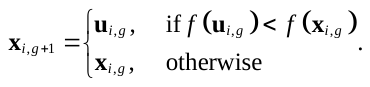

To address sensitivity to control parameters in traditional DE, JADE incorporates a dynamic parameter adjustment mechanism. Mutation factor
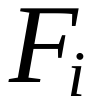
 and crossover probability
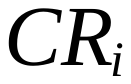
 are dynamically updated based on the evolutionary feedback. The
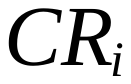
 value is drawn from a normal distribution (
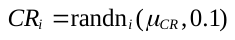
). The mean
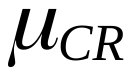
 is updated as:

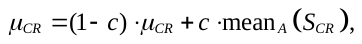

Here,
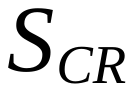
 is the set of successful
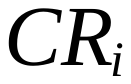
 values from the previous generation. Similarly,
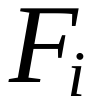
 is sampled from a Cauchy distribution with mean
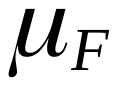
 (
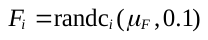
). The
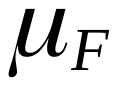
 is updated using the Lehmer mean:

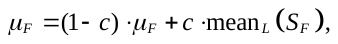

where
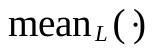
 represents the Lehmer mean:

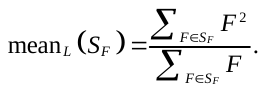

The Lehmer mean promotes larger mutation factors, improving search efficiency. Both
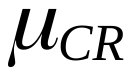
 and
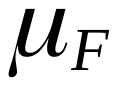
 are initialized to 0.5. Besides,
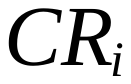
 and
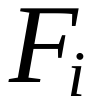
 truncated within
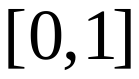
.

JADE addresses the tedious parameter tuning in traditional DE and introduces two additional parameters,
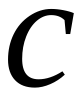
 (adaptive rate) and
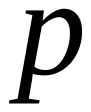
 (greediness of mutation strategy). Experimental results suggest robust default values for various problems:
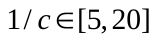
 and
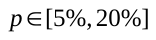
.

### Platforms

This study was implemented using Python for model reading, operations, and genetic algorithm execution, with PyCharm as the integrated development environment (IDE) for efficient development and debugging. Experiments were conducted on a Windows system with an Intel(R) Core(TM) i9-14900HX processor.

The COBRApy (Ebrahim et al., 2013) library was used for constraint-based metabolic network reconstruction and analysis. It provides comprehensive functionality for constructing, modifying, and simulating metabolic models. Additionally, the StrainDesign (Schneider et al., 2022) toolbox, which integrates optimization algorithms and pathway visualization tools, supported the strain design tasks. For model optimization, Gurobi 11.0 (Gurobi Optimization), a high-performance mathematical programming solver, was utilized to solve linear and quadratic programming problems. Its capabilities significantly enhanced the efficiency of genetic algorithms, especially during large-scale gene editing strategy searches.

### Supplementary tables

**Table S1 Details of the ecYeast 8.3.4 and eciML1515**

| Content | ecYeast 8.3.4 | eciML1515 |
| --- | --- | --- |
| All reaction | 8144 | 6085 |
| Metabolite | 4180 | 3594 |
| Gene | 1148 | 1505 |
| Medium | 16 | 26 |
| Boundary | 487 | 675 |
| Demands | 2 | 12 |
| Exchange reaction | 484 | 662 |
| Gene that encode enzyme | 968 | 1259 |
| Enzyme synthesis reaction | 968 | 1259 |
| Enzyme with abundance data | 968 | 1259 |
| Arm reaction | 520 | 456 |
| Metabolic reaction | 6655 | 4369 |
| Protein pool | 1 | 1 |
| cell compartment | 14 | 3 |

**Table S2** **Strains and plasmids used in this study**

| Strain | Description | Origin |
| --- | --- | --- |
| *E. coli* DH5α | Competent cells | Sangon Biotech, Shanghai, China |
| *E. coli* MG1655 | Wide-type strain | Lab stock |
| MGHT-10-1 | *E. coli* MG1655, trpRΔ, *tnaA*Δ, *pheA*Δ, *tyrA*Δ, P*_J23101_*-*trpCBA,* P*_J23101_*-*trpE*D* | Lab stock |
| MGT-1 | MGHT-10-1, *pgi*Δ | This study |
| MGT-2 | MGHT-10-1, *fsaA*Δ | This study |
| MGT-3 | MGHT-10-1, *rpe*Δ | This study |
| MGT-4 | MGHT-10-1, P*_J23101_*-*aroL* | This study |
| MGT-5 | MGHT-10-1, P*_J23101_*-*aroC* | This study |
| MGT-6 | MGHT-10-1, *zwf*Δ | This study |
| MGT-7 | MGHT-10-1, *pgl*Δ | This study |
| MGT-8 | MGHT-10-1, *fsaB*Δ | This study |
| MGT-9 | MGHT-10-1, P*_J23101_*-*aroF* | This study |
| MGT-10 | MGHT-10-1, P*_J23101_*-*talA* | This study |
| MGT-11 | MGHT-10-1, P*_J23114_*-*tpiA* | This study |
| MGT-12 | MGT-1, P*_J23101_*-*gnd* | This study |
| MGT-13 | MGT-2, *fsaB*Δ | This study |
| MGT-14 | MGT-12, P*_J23114_*-*lpd* | This study |
| MGT-15 | MGT-13, P*_J23114_*-*lpd* | This study |
| MGT-16 | MGT-14, P*_J23114_*-*rpe* | This study |
| MGT-17 | MGT-15, P*_J23101_*-*aroC* | This study |
| MGT-18 | MGT-15, P*_J23101_*-*aroL* | This study |
| MGT-19 | MGT-15, *dhaK*Δ | This study |
| MGT-20 | MGT-16, P*_J23101_*-*aroG** | This study |
| Plasmids |  | This study |
| pEcCas | pCas, *sacB*, P_rhaB_-sgRNA-pMB1 | Lab stock |
| pEcgRNA | pTargetF, *ccdB* | Lab stock |
| pEcgRNA-*pgi* | pEcgRNA, target *pgi* in MGHT-10-1 | This study |
| pEcgRNA-*fsaA* | pEcgRNA, target *fsaA* in MGHT-10-1 | This study |
| pEcgRNA-*rpe* | pEcgRNA, target *rpe* in MGHT-10-1 | This study |
| pEcgRNA-*yncI* | pEcgRNA, target *ycnI* for overexpressing *aroL*/*aroC*/*aroF*/*talA* | This study |
| pEcgRNA-*zwf* | pEcgRNA, target *zwf* in MGHT-10-1 | This study |
| pEcgRNA-*pgl* | pEcgRNA, target *pgl* in MGHT-10-1 | This study |
| pEcgRNA-*fsaB* | pEcgRNA, target *fsaB* in MGHT-10-1 | This study |
| pEcgRNA-*tpiA* | pEcgRNA, target *tpiA* in MGHT-10-1 | This study |
| pEcgRNA-*gnd* | pEcgRNA, target *gnd* for overexpressing *gnd* | This study |
| pEcgRNA-*lpd* | pEcgRNA, target *lpd* for knocking down *lpd* | This study |
| pEcgRNA-*aroG** | pEcgRNA, target *yjiV* for overexpressing *aroG** | This study |

Note：*trpE** indicates that the 40th amino acid in TrpE from serine (S) to phenylalanine (F), aroG* indicates that the180th amino acid in AroG from serine (S) to phenylalanine (F)

**Table S3 The primer sequences used in this study**

| Primer | Sequences (5’- 3’) |
| --- | --- |
| *pgi-*sgRNA-F | ttctgacctcggcccatacagttttagagctagaaatagcaag |
| *pgi-*sgRNA-R | tgtatgggccgaggtcagaaactagtattatacctaggactgagctag |
| *pgi-*U-F | tggcacaagggaagagcgg |
| *pgi-*U-R | gaaacaaagtgcatgttcaggtggagcggtctgcgttggattgatg |
| *pgi-*D-F | ccacctgaacatgcactttgtttc |
| *pgi-*D-R | acggtgcatatactggtcatacg |
| *fsaA-*sgRNA-F | aagtttgagcaggactggcagttttagagctagaaatagcaag |
| *fsaA-*sgRNA-R | tgccagtcctgctcaaacttactagtattatacctaggactgagctag |
| *fsaA*-U-F | acggtttatcgcgcagaacg |
| *fsaA*-U-R | accgcaacaacgtctgaagtatcc |
| *fsaA*-D-F | ggatacttcagacgttgttgcggtaccggacgttctgcatcctc |
| *fsaA*-D-R | acatatcgcgattacgccag |
| *rpe*-sgRNA-F | ctttaatcagttgcagcgtggttttagagctagaaatagcaag |
| *rpe*-sgRNA-R | cacgctgcaactgattaaagactagtattatacctaggactgagctag |
| *rpe*-U-F | tgcacatcatcaccaccaatcac |
| *rpe*-U-R | ctcaaggagaagcggatgaaacaatgcgcagtgaactggcaaag |
| *rpe*-D-F | tgtttcatccgcttctccttgag |
| *rpe*-D-R | accgctacggttacaacggc |
| *ycnI-*sgRNA-F | ttccttgttgtcgtatatcagttttagagctagaaatagcaag |
| *ycnI-*sgRNA-R | tgatatacgacaacaaggaaactagtattatacctaggactgagctag |
| *aroL*-U-F | tcaaacagaaaggccatggacaccaaagagcgttgagatatatc |
| *aroL*-U-R | agagctagcataatacctaggactgagctagctgtaaaactttgaatactcacctggcg |
| *aroL-*F | ggtattatgctagctctagagaaagaggagaaatactagatgacacaacctctttttctgatc |
| *aroL-*R | aaggcccagtctttcgactgagcctttcgttttatttgtcaacaattgatcgtctgtgcc |
| *aroL-*D-F | tcgaaagactgggcctttcgttttatctgttgtttgtcggtgaacgctctcctgagtaggacaaattgtttctcaccgtatgtgcag |
| *aroL-*D-R | cccgtagaatccatgaggatcctcgaatgcatgatgtaatttcc |
| *aroC-*F | ggtattatgctagctctagagaaagaggagaaatactagatggctggaaacacaattgg |
| *aroC-*R | aaggcccagtctttcgactgagcctttcgttttatttgttaccagcgtggaatatcagtc |
| *zwf-*sgRNA-F | gcgccgaaaccgtatcaggcgttttagagctagaaatagcaag |
| *zwf-*sgRNA-R | gcctgatacggtttcggcgcactagtattatacctaggactgagctag |
| *zwf-*U-F | tggatagtgttcataaggctggtgcg |
| *zwf-*U-R | tggtcgttcctggaatgagtttgag |
| *zwf-*D-F | ctcaaactcattccaggaacgaccatcctcgaagtattcgccaacctg |
| *zwf-*D-R | aacttaaggagaatgacatggcgg |
| *pgl-*sgRNA-F | tatgaatatcagccgcccaagttttagagctagaaatagcaag |
| *pgl-*sgRNA-R | ttgggcggctgatattcataactagtattatacctaggactgagctag |
| *pgl-*U-F | agttagatgtgcgaatgacatgcc |
| *pgl-*U-R | gtaaatcagcggttagtgtgcgttcatgaatgctcctttgcatttagc |

**Table S3 The primer sequences used in this study (continued)**

| Primer | Sequences (5’- 3’) |
| --- | --- |
| *pgl-*D-F | acgcacactaaccgctgatttac |
| *pgl-*D-R | actggtcaacgagccgttcg |
| *fsaB*-sgRNA-F | acgcggcagcacttcccatagttttagagctagaaatagcaag |
| *fsaB*-sgRNA-R | tatgggaagtgctgccgcgtactagtattatacctaggactgagctag |
| *fsaB-*U-F | atacccgccgctaacagacg |
| *fsaB-*U-R | aggagcaattatggaccgcattattc |
| *fsaB-*D-F | gaataatgcggtccataattgctccttgccagacgttcgacttctgc |
| *fsaB-*D-R | aacacctgctggcgcaattc |
| *aroF-*F | ggtattatgctagctctagagaaagaggagaaatactagatgcaaaaagacgcgctgaa |
| *aroF-*R | gaaaggcccagtctttcgactgagcctttcgttttatttgttaagccacgcgagccgtc |
| *talA-*F | ggtattatgctagctctagagaaagaggagaaatactagatgaacgagttagacggcat |
| *talA-*R | aaggcccagtctttcgactgagcctttcgttttatttgttatagtttggcggcaagaag |
| *tpiA-*U-F | tgagccggagttgcagatttgc |
| *tpiA-*U*-*R | gtcctaggtacaatgctagctctagagaaagaggggacaaactagatgcgacatcctttagtgatgg |
| *tpiA-*D-F | ctctagagctagcattgtacctaggactgagctagccataaatgtcatttattcaaaccttcaagcg |
| *tpiA-*D*-*R | aacaccagatacggcaacgac |
| *gnd*-sgRNA-F | atcaccgcgctgaatgctcggttttagagctagaaatagcaag |
| *gnd*-sgRNA-R | cgagcattcagcgcggtgatactagtattatacctaggactgagctag |
| *gnd-*U-F | atcggcaccaatataggtaacgc |
| *gnd-*U*-*R | AGTCCTAGGTATTATGCTAGCTCTAGAGAAAGAGGAGAAATACTAGatgtccaagcaacagatcggc |
| *gnd-*D-F | TCTCTAGAGCTAGCATAATACCTAGGACTGAGCTAGCTGTAAAtggaatgttcgcaaataagtatacaaagtac |
| *gnd-*D*-*R | agtatctattctgatacggttgttgattgg |
| *lpd*-sgRNA-F | taagtaagtgactggggtgagttttagagctagaaatagcaag |
| *lpd*-sgRNA-R | tcaccccagtcacttacttaactagtattatacctaggactgagctag |
| *lpd-*U-F | aggcatcatcgagctgtctcg |
| *lpd-*U-R | tctctagagctagcattgtacctaggactgagctagccataaatgctgcacccagaaatccatag |
| *lpd-*D-F | tcctaggtacaatgctagctctagagaaagaggggacaaactagatgagtactgaaatcaaaactcaggtcg |
| *lpd-*D-R | tcagttccagcgcgtcagtg |
| *rpe*-U1-F | aatgcgcgctccatcagaac |
| *rpe-*U1*-*R | aaccctacagctaatcaaggaaaatggctgtaaagcgggtc |
| *rpe-*D1-F | ccattttccttgattagctgtagggttcggtcaacatgctcggagg |
| *rpe-*D1-R | gtcctaggtacaatgctagctctagagaaagaggggacaaactagagcggatgaaacagtatttgattgc |

**Table S3 The primer sequences used in this study (continued)**

| Primer | Sequences (5’- 3’) |
| --- | --- |
| *rpe-*D2-F | ctctagagctagcattgtacctaggactgagctagccataaatctccttgagaattattttttcgcggg |
| *rpe-*D2-R | agcgcatctggcggagatc |
| *dhaK-sgRNA-F* | caatcacccgatcgccagacgttttagagctagaaatagcaag |
| *dhaK-sgRNA-R* | gtctggcgatcgggtgattgactagtattatacctaggactgagctag |
| *dhaK-*U-F | actgattacgccgtccgcg |
| *dhaK-*U-R | gtgcaagacgtactggacgaacataaccgcctgaccacacgttg |
| *dhaK-*D-F | tgttcgtccagtacgtcttgcac |
| *dhaK-*D-R | tacggctgagaatgcaggcg |
| *yjiV-*sgRNA-F | ccatgactggatgagcataggttttagagctagaaatagcaag |
| *yjiV-*sgRNA-R | ctatgctcatccagtcatggactagtattatacctaggactgagctag |
| *aroG-*U-F | actatcgtactgccgagcactc |
| *aroG-*U-R | aggactgagctagctgtaaaatccagcgtcacttcagcg |
| *aroG-*F | tttacagctagctcagtcctagg |
| *aroG-*R | ttacccgcgacgcgcttttac |
| *aroG-*D-F | gtaaaagcgcgtcgcgggtaacaaataaaacgaaaggctcagtcgaaag |
| *aroG-*D-R | tcgttaaagcctgaaaatggcg |

**Table S4 Experimental results of different searching algorithms**

| Algorithm | 2-phenylethanol | spermidine | L-tryptophan |
| --- | --- | --- | --- |
| binGA (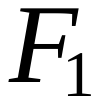)  (Std) | 1.41E-01  (5.58E-05) | \ | \ |
| binGA (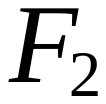)  (Std) | 1.46E-01  (2.00E-04) | \ | \ |
| symGA (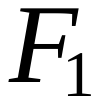)  (Std) | 1.62E-01  (6.06E-03) | \ | \ |
| symGA (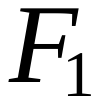, non-elitism)  (Std) | 1.64E-01  (4.87E-03) | \ | \ |
| JADE  (Std) | 1.71E-01  (2.00E-04) | 1.89E-01  (5.05E-03) | 4.49E-02  (2.30E-03) |

**Table S5 The ranked strategies obtained by JADE in *S. cerevisiae***

| Product | Individual  (Strategy) | 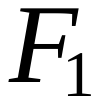 | 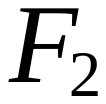 | 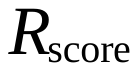 | Number of targets | Number of exp targets |
| --- | --- | --- | --- | --- | --- | --- |
| 2-PE | 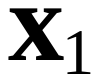 | 0.1707 | 0.1748 | 0.1778 | 11 | 3 |
|  | 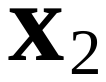 | 0.1711 | 0.1748 | 0.1768 | 15 | 2 |
|  | 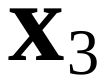 | 0.1705 | 0.1745 | 0.1765 | 12 | 2 |
|  | 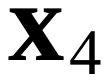 | 0.1707 | 0.1750 | 0.1760 | 9 | 1 |
|  | 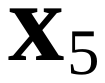 | 0.1705 | 0.1748 | 0.1758 | 9 | 1 |
|  | 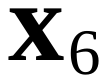 | 0.1705 | 0.1748 | 0.1758 | 9 | 1 |
|  | 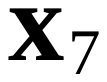 | 0.1705 | 0.1748 | 0.1758 | 9 | 1 |
|  | 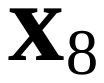 | 0.1706 | 0.1746 | 0.1756 | 12 | 1 |
|  | 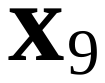 | 0.1705 | 0.1746 | 0.1756 | 11 | 1 |
|  | 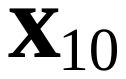 | 0.1705 | 0.1745 | 0.1755 | 12 | 1 |
| Spermidine | 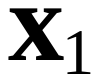 | 0.1934 | 0.2001 | 0.2021 | 19 | 2 |
|  | 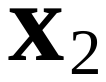 | 0.1918 | 0.1990 | 0.2010 | 14 | 2 |
|  | 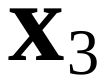 | 0.1918 | 0.1989 | 0.2009 | 15 | 2 |
|  | 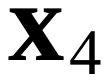 | 0.1919 | 0.1989 | 0.2009 | 16 | 2 |
|  | 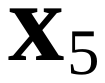 | 0.1905 | 0.1974 | 0.1994 | 17 | 2 |
|  | 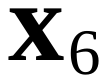 | 0.1910 | 0.1982 | 0.1992 | 14 | 1 |
|  | 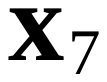 | 0.1906 | 0.1972 | 0.1982 | 20 | 1 |
|  |  | 0.1853 | 0.1920 | 0.1950 | 19 | 3 |
|  |  | 0.1829 | 0.1899 | 0.1909 | 16 | 1 |
|  |  | 0.1806 | 0.1876 | 0.1886 | 16 | 1 |

**Table S6 Experimentally validated targets for 2-PE and spermidine**

| Product | Gene ID | Short name | Action | Source |
| --- | --- | --- | --- | --- |
| 2-PE | YGL148W | *ARO2* | OE | 10.1101/2023.01.31.526512 |
|  | YNL316C | *PHA2* | OE | 10.1101/2023.01.31.526512 |
|  | YDR380W | *ARO10* | OE | 10.1101/2023.01.31.526512 |
|  | YDR127W | *ARO1* | OE | 10.1101/2023.01.31.526512 |
|  | YBR249C | *ARO4* | OE | 10.1101/2023.01.31.526512 |
|  | YPR060C | *ARO7* | OE | 10.1101/2023.01.31.526512 |
|  | YNL241C | *ZWF1* | OE | 10.1101/2023.01.31.526512 |
|  | **YGL202W** | *ARO8* | OE | 10.12211/2096-8280.2020-096 |
|  | \ | *MT2* | OE | 10.12211/2096-8280.2020-096 |
|  | YPR074C | *TKL1* | OE | 10.12211/2096-8280.2020-096 |
|  | **YAL038W** | *CDC19* | OE | 10.12211/2096-8280.2020-096 |
|  | **YDR035W** | *ARO3* | KD | 10.12211/2096-8280.2020-096 |
|  | **YOL052C** | *SPE2* | OE | 10.1038/s41929-021-00631-z |
| spermidine | **YPR069C** | *SPE3* | OE | 10.1038/s41929-021-00631-z |
|  | **YDR502C** | *SAM2* | OE | 10.1038/s41929-021-00631-z |
|  | YJR148W | *BAT2* | OE | 10.1038/s41929-021-00631-z |
|  | **YLR017W** | *MEU1* | OE | 10.1038/s41929-021-00631-z |
|  | **YML022W** | *APT1* | OE | 10.1038/s41929-021-00631-z |
|  | YLL028W | *TPO1* | OE | 10.1038/s41929-021-00631-z |
|  | **YOL061W** | *PRS5* | OE | 10.1038/s41929-021-00631-z |
|  | **YKL184W** | *ODC(SPE1)* | OE | 10.1038/s41929-021-00631-z |
|  | YOR130C | *ORT1* | OE | 10.1038/s41929-021-00631-z |
|  | YPR021C | *AGC1* | OE | 10.1038/s41929-021-00631-z |
|  | **YOR375C** | *GDH1* | OE | 10.1038/s41929-021-00631-z |
|  | YLR146C | *SPE4* | KO | 10.1038/s41929-021-00631-z |
|  | YMR020W | *FMS1* | KO | 10.1038/s41929-021-00631-z |
|  | YNL141W | *AAH1* | KO | 10.1038/s41929-021-00631-z |
|  | YPL052W | *OAZ1* | KO | 10.1038/s41929-021-00631-z |
|  | **YLR438W** | *CAR2* | KO | 10.1038/s41929-021-00631-z |
|  | YJL088W | *ARG3* | KD | 10.1038/s41929-021-00631-z |
|  | YER037W | *PHM8* | KO | 10.1038/s41929-021-00631-z |
|  | YJR139C | *HOM6* | KD | 10.1038/s41929-021-00631-z |
|  | YDR481C | *PHO8* | KD | 10.1038/s41929-021-00631-z |

Note: Genes shown in bold indicate those were also predicted by MetaStrain.

**Table S7 Results of ingle-target redundancy analysis**

| Strategy ID | Original edits | Minimal edits | Reduced | Original fitness | Minimal fitness | Fitness ratio |
| --- | --- | --- | --- | --- | --- | --- |
| 1 | 10 | 5 | 5 | 0.0431 | 0.0424 | 0.9833 |
| 2 | 8 | 6 | 2 | 0.0500 | 0.0499 | 0.9971 |
| 3 | 8 | 6 | 2 | 0.0447 | 0.0001 | 0.0022 |
| 4 | 11 | 5 | 6 | 0.0442 | 0.0424 | 0.9589 |
| 5 | 11 | 6 | 5 | 0.0436 | 0.0441 | 1.0131 |
| 6 | 6 | 6 | 0 | 0.0422 | 0.0422 | 1.0000 |
| 7 | 14 | 4 | 10 | 0.0447 | 0.0001 | 0.0022 |
| 8 | 9 | 6 | 3 | 0.0482 | 0.0493 | 1.0225 |
| 9 | 10 | 5 | 5 | 0.0414 | 0.0001 | 0.0024 |
| 10 | 9 | 6 | 3 | 0.0458 | 0.0458 | 1.0000 |
| 11 | 10 | 6 | 4 | 0.0486 | 0.0470 | 0.9662 |
| 12 | 6 | 5 | 1 | 0.0496 | 0.0493 | 0.9943 |
| 13 | 9 | 4 | 5 | 0.0442 | 0.0256 | 0.5788 |
| 14 | 10 | 5 | 5 | 0.0424 | 0.0446 | 1.0522 |
| 15 | 10 | 7 | 3 | 0.0454 | 0.0001 | 0.0022 |
| 16 | 13 | 4 | 9 | 0.0474 | 0.0136 | 0.2863 |
| 17 | 7 | 5 | 2 | 0.0419 | 0.0001 | 0.0024 |
| 18 | 7 | 5 | 2 | 0.0416 | 0.0001 | 0.0024 |
| 19 | 7 | 6 | 1 | 0.0441 | 0.0441 | 1.0000 |
| 20 | 8 | 5 | 3 | 0.0445 | 0.0445 | 1.0000 |

**Table S8 Top 30 4-target combinatorial strategies**

| Strategy ID |  |  | Combination genes targets |  | Percentage |
| --- | --- | --- | --- | --- | --- |
| 14 | 10 | 0.042372 | b0116(KD) + b0825(KO) + b1200(KO) + b3946(KO) | 0.040656 | 95.95 |
| 2 | 8 | 0.050048 | b4025(KO) + b0116(KD) + b2029(OE) + b3386(KD) | 0.040021 | 79.96 |
| 10 | 9 | 0.045764 | b0388(OE) + b0116(KD) + b0825(KO) + b3946(KO) | 0.039669 | 86.68 |
| 19 | 7 | 0.044145 | b0388(OE) + b0116(KD) + b0825(KO) + b3946(KO) | 0.039669 | 89.86 |
| 14 | 10 | 0.042372 | b2329(OE) + b0116(KD) + b0825(KO) + b3946(KO) | 0.039669 | 93.62 |
| 5 | 11 | 0.043576 | b1693(OE) + b0116(KD) + b0825(KO) + b3946(KO) | 0.039669 | 91.03 |
| 11 | 10 | 0.048623 | b1693(OE) + b0116(KD) + b0825(KO) + b3946(KO) | 0.039669 | 81.58 |
| 8 | 9 | 0.048247 | b4025(KO) + b0388(OE) + b0116(KD) + b3386(KO) | 0.038890 | 80.61 |
| 20 | 8 | 0.044467 | b4025(KO) + b0388(OE) + b0116(KD) + b3386(KO) | 0.038890 | 87.46 |
| 12 | 6 | 0.049622 | b4025(KO) + b0754(OE) + b0116(KD) + b3386(KO) | 0.038890 | 78.37 |
| 2 | 8 | 0.050048 | b4025(KO) + b0754(OE) + b0116(KD) + b3386(KD) | 0.038760 | 77.45 |
| 13 | 9 | 0.044226 | b4025(KO) + b0754(OE) + b0116(KD) + b3386(KD) | 0.038760 | 87.64 |
| 9 | 10 | 0.041380 | b4025(KO) + b1693(OE) + b0116(KD) + b3386(KD) | 0.038760 | 93.67 |
| 18 | 7 | 0.041565 | b0388(OE) + b4094(OE) + b0116(KD) + b1198(KO) | 0.038447 | 92.50 |
| 1 | 10 | 0.043073 | b2329(OE) + b4094(OE) + b0116(KD) + b1198(KO) | 0.038447 | 89.26 |
| 16 | 13 | 0.047429 | b2329(OE) + b4094(OE) + b0116(KD) + b1198(KO) | 0.038447 | 81.06 |
| 4 | 11 | 0.044170 | b1693(OE) + b4094(OE) + b0116(KD) + b1200(KO) | 0.038447 | 87.04 |
| 16 | 13 | 0.047429 | b1693(OE) + b4094(OE) + b0116(KD) + b1198(KO) | 0.038447 | 81.06 |
| 18 | 7 | 0.041565 | b0388(OE) + b0116(KD) + b2029(OE) + b1198(KO) | 0.037112 | 89.29 |
| 7 | 14 | 0.044673 | b4025(KD) + b0116(KD) + b2029(OE) + b3386(KO) | 0.037004 | 82.83 |
| 7 | 14 | 0.044673 | b0116(KD) + b2029(OE) + b3386(KO) + b1199(KD) | 0.036621 | 81.98 |
| 12 | 6 | 0.049622 | b0116(KD) + b2029(OE) + b3386(KO) + b1200(KD) | 0.036621 | 73.80 |

| 7 | 14 | 0.044673 | b0754(OE) + b4094(OE) + b0116(KD) + b1199(KD) | 0.035773 | 80.08 |
| --- | --- | --- | --- | --- | --- |
| 15 | 10 | 0.045420 | b0388(OE) + b4094(OE) + b0116(KD) + b1199(KD) | 0.035773 | 78.76 |
| 3 | 8 | 0.044685 | b4094(OE) + b0116(KD) + b2029(OE) + b3386(KD) | 0.035610 | 79.69 |
| 7 | 14 | 0.044673 | b4094(OE) + b0116(KD) + b2029(OE) + b3386(KO) | 0.035520 | 79.51 |
| 11 | 10 | 0.048623 | b1693(OE) + b3919(KD) + b0116(KD) + b0825(KO) | 0.035264 | 72.52 |
| 14 | 10 | 0.042372 | b2329(OE) + b3919(KD) + b0116(KD) + b0825(KO) | 0.035264 | 83.22 |
| 3 | 8 | 0.044685 | b0116(KD) + b2029(OE) + b3386(KD) + b0825(KO) | 0.035231 | 78.84 |
| 4 | 11 | 0.044170 | b4094(OE) + b0116(KD) + b1033(KO) + b0825(KO) | 0.035162 | 79.61 |

**Table S9 Top 30 5-target combinatorial strategies**

| Strategy ID |  |  | Combination genes targets |  | Percentage |
| --- | --- | --- | --- | --- | --- |

| 2 | 8 | 0.050048 | b4025(KO) + b0754(OE) + b0116(KD) + b2029(OE) + b3386(KD) | 0.049832 | 99.57 |
| --- | --- | --- | --- | --- | --- |

| 8 | 9 | 0.048247 | b4025(KO) + b0388(OE) + b0116(KD) + b2029(OE) + b3386(KO) | 0.049340 | 102.27 |
| --- | --- | --- | --- | --- | --- |
| 12 | 6 | 0.049622 | b4025(KO) + b0754(OE) + b0116(KD) + b2029(OE) + b3386(KO) | 0.049340 | 99.43 |
| 14 | 10 | 0.042372 | b2329(OE) + b3919(KD) + b0116(KD) + b0825(KO) + b3946(KO) | 0.044583 | 105.22 |
| 11 | 10 | 0.048623 | b1693(OE) + b3919(KD) + b0116(KD) + b0825(KO) + b3946(KO) | 0.044583 | 91.69 |
| 20 | 8 | 0.044467 | b4025(KO) + b0388(OE) + b4094(OE) + b0116(KD) + b3386(KO) | 0.044467 | 100.00 |
| 8 | 9 | 0.048247 | b0388(OE) + b3919(KD) + b0116(KD) + b2029(OE) + b3386(KO) | 0.042752 | 88.61 |
| 1 | 10 | 0.043073 | b4025(KO) + b2329(OE) + b4094(OE) + b0116(KD) + b1198(KO) | 0.042355 | 98.33 |
| 16 | 13 | 0.047429 | b4025(KO) + b2329(OE) + b4094(OE) + b0116(KD) + b1198(KO) | 0.042355 | 89.30 |
| 4 | 11 | 0.044170 | b4025(KO) + b1693(OE) + b4094(OE) + b0116(KD) + b1200(KO) | 0.042355 | 95.89 |
| 16 | 13 | 0.047429 | b4025(KO) + b1693(OE) + b4094(OE) + b0116(KD) + b1198(KO) | 0.042355 | 89.30 |
| 10 | 9 | 0.045764 | b0388(OE) + b4094(OE) + b0116(KD) + b0825(KO) + b3946(KO) | 0.041934 | 91.63 |
| 8 | 9 | 0.048247 | b4025(KO) + b0388(OE) + b3919(KD) + b0116(KD) + b3386(KO) | 0.041816 | 86.67 |
| 19 | 7 | 0.044145 | b0388(OE) + b2744(KO) + b0116(KD) + b0825(KO) + b3946(KO) | 0.041671 | 94.40 |
| 5 | 11 | 0.043576 | b1693(OE) + b2744(KO) + b0116(KD) + b0825(KO) + b3946(KO) | 0.041671 | 95.63 |
| 7 | 14 | 0.044673 | b4025(KD) + b0116(KD) + b2029(OE) + b3386(KO) + b1199(KD) | 0.041148 | 92.11 |
| 7 | 14 | 0.044673 | b4025(KD) + b0754(OE) + b0116(KD) + b2029(OE) + b3386(KO) | 0.041141 | 92.09 |
| 14 | 10 | 0.042372 | b0116(KD) + b3125(KO) + b0825(KO) + b1200(KO) + b3946(KO) | 0.040669 | 95.98 |
| 14 | 10 | 0.042372 | b2925(OE) + b0116(KD) + b0825(KO) + b1200(KO) + b3946(KO) | 0.040656 | 95.95 |
| 14 | 10 | 0.042372 | b0116(KD) + b3553(KO) + b0825(KO) + b1200(KO) + b3946(KO) | 0.040656 | 95.95 |
| 14 | 10 | 0.042372 | b4383(OE) + b0116(KD) + b0825(KO) + b1200(KO) + b3946(KO) | 0.040605 | 95.83 |
| 3 | 8 | 0.044685 | b0754(OE) + b4094(OE) + b0116(KD) + b2029(OE) + b3386(KD) | 0.040432 | 90.48 |
| 13 | 9 | 0.044226 | b4025(KO) + b0754(OE) + b2744(KO) + b0116(KD) + b3386(KD) | 0.040297 | 91.12 |
| 11 | 10 | 0.048623 | b1693(OE) + b3919(KD) + b0116(KD) + b2029(OE) + b0825(KO) | 0.040241 | 82.76 |
| 2 | 8 | 0.050048 | b4025(KO) + b0116(KD) + b2029(OE) + b3386(KD) + b0825(KD) | 0.040195 | 80.31 |
| 19 | 7 | 0.044145 | b0388(OE) + b4226(KO) + b0116(KD) + b0825(KO) + b3946(KO) | 0.040163 | 90.98 |
| 5 | 11 | 0.043576 | b1693(OE) + b4226(KO) + b0116(KD) + b0825(KO) + b3946(KO) | 0.040163 | 92.17 |
| 11 | 10 | 0.048623 | b1693(OE) + b4226(KO) + b0116(KD) + b0825(KO) + b3946(KO) | 0.040163 | 82.60 |
| 7 | 14 | 0.044673 | b4025(KD) + b4094(OE) + b0116(KD) + b2029(OE) + b3386(KO) | 0.040025 | 89.60 |
| 7 | 14 | 0.044673 | b0754(OE) + b4094(OE) + b0116(KD) + b2029(OE) + b3386(KO) | 0.039841 | 89.18 |

### Supplementary figures

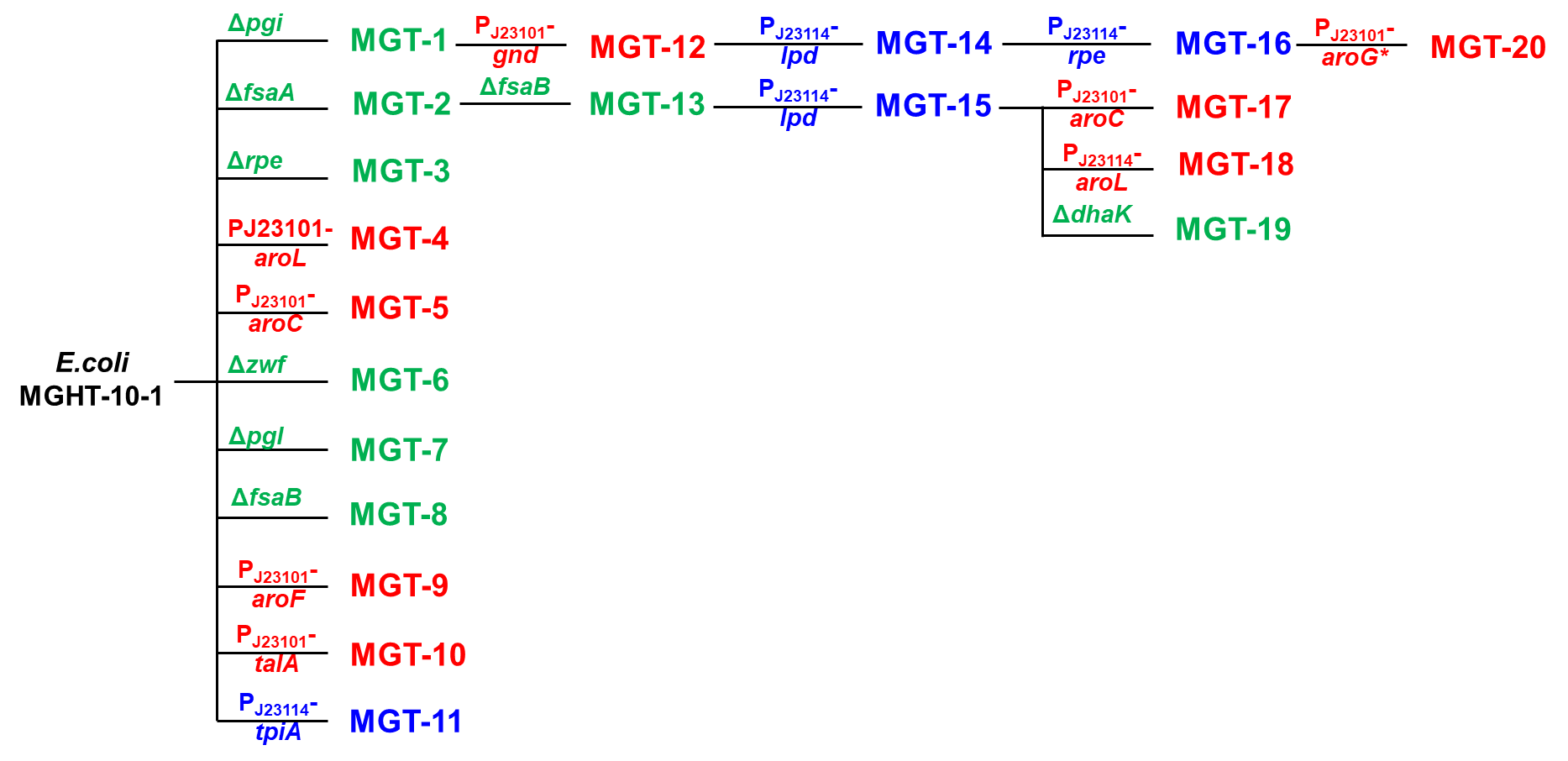

**Fig. S1 Experimental design technical route.** The experimental plans were designed layer by layer based on the predicted combination targets and metabolic pathways. Green represents knockout, red represents overexpression, and blue represents knockdown.

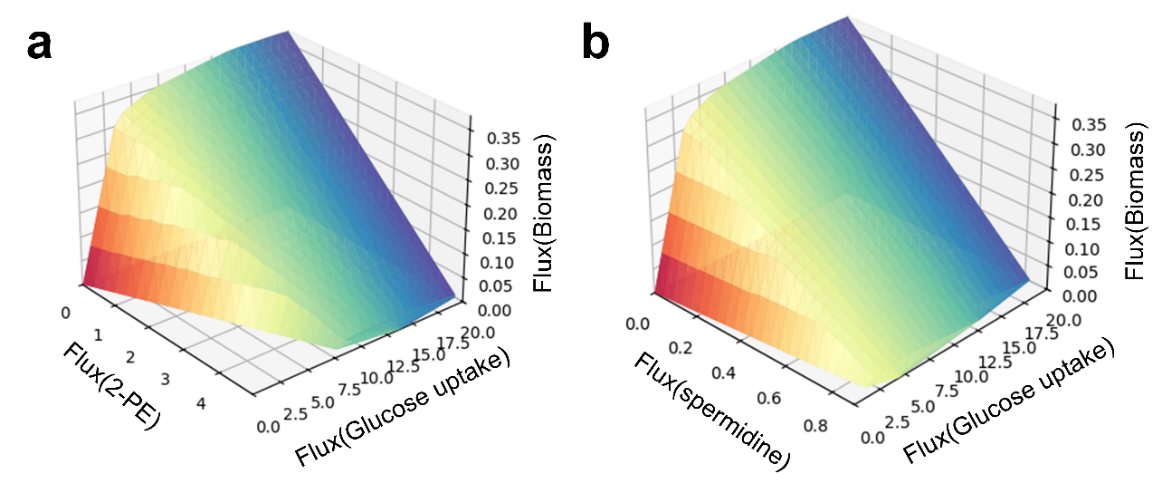

**Fig. S2 Phase plane of *S. cerevisiae*.** (a, b) A phenotypic phase plane analysis for 2-PE (a) and spermidine (b) production in the wild-type *S. cerevisiae* model, illustrating the inherent metabolic trade-offs.

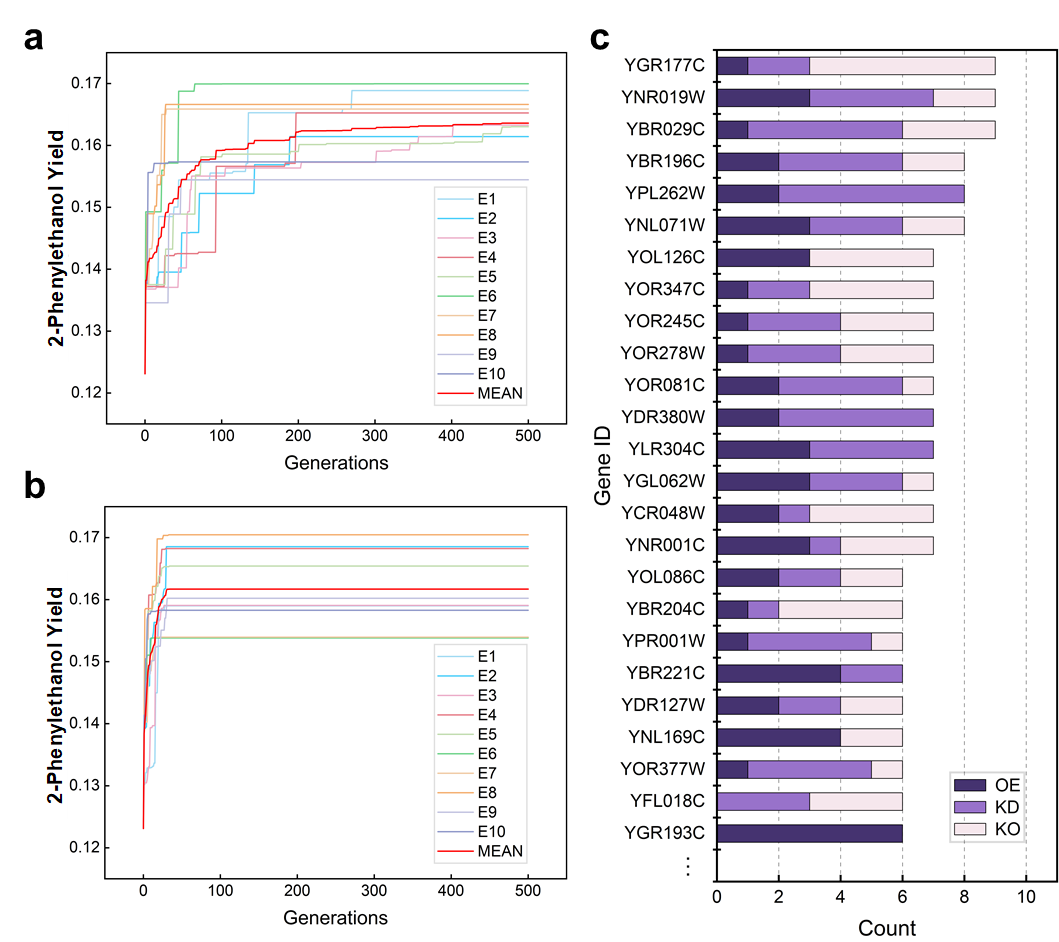

**Fig. S3 Searching results of symGA.** (a, b) The convergence curve of the symGA with (b) and without (a) an elite retention strategy searching for combined targets for overexpression of 2-PE in *S. cerevisiae*. (c) Accumulative histograms summarizing the frequency of each gene's appearance in the optimal solutions identified across ten independent runs.

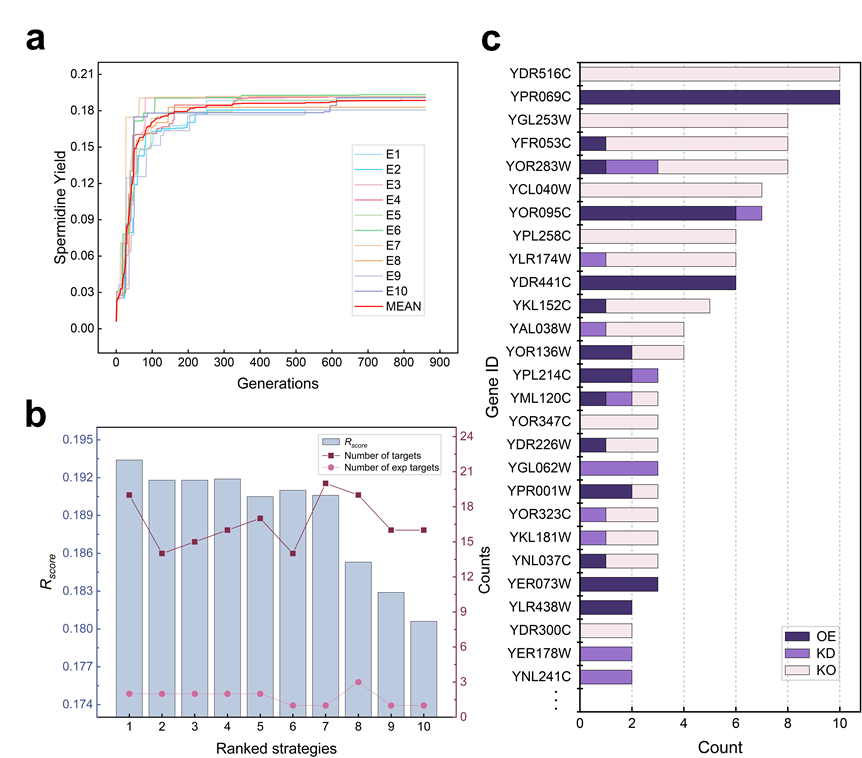

**Fig. S4** **Results of JADE using spermidine as the product.** (a) The convergence profile of the JADE algorithm during the search for optimal combinational gene-editing strategies. (b) Detail information of optimal strategies predicted by JADE, including the comprehensive scores, number of targets and validated targets of each strategy. (c) An accumulative histogram summarizing the top-performing gene targets identified across ten independent JADE runs.

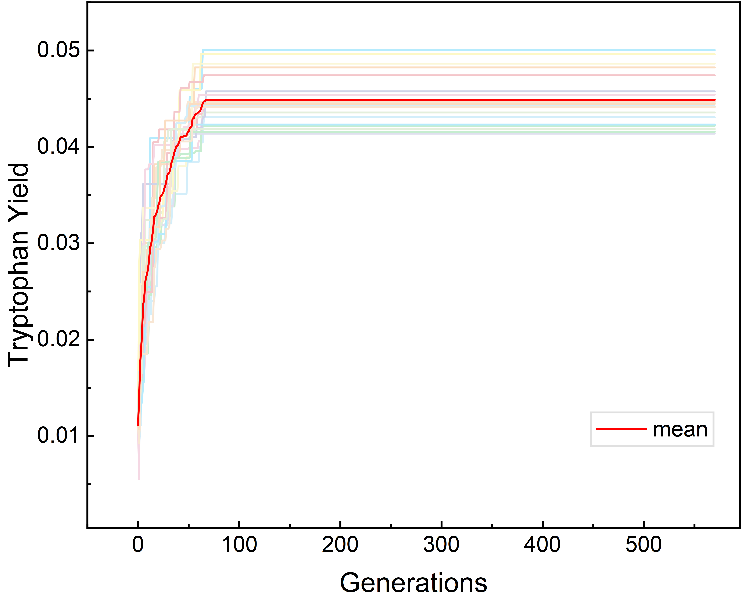

**Fig. S5 The search process of JADE**. Convergent curves of twenty repeated experiments for identifying combined targets for excessive expression of tryptophan in *E. coli*.

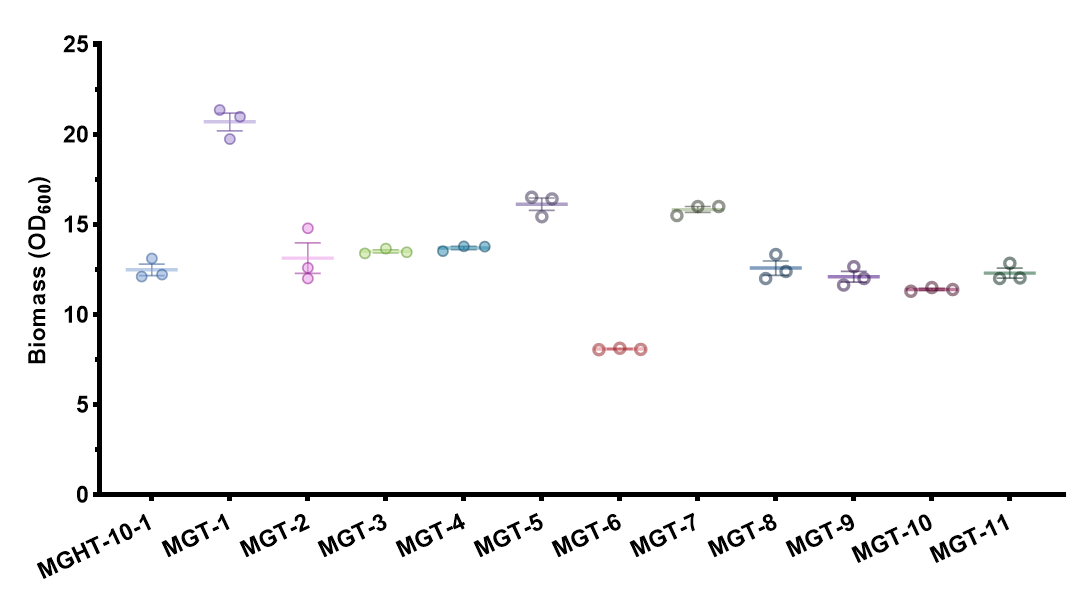

**Fig. S6 The biomass of the recombinant strain.**

### Supplementary references

Domenzain, I., Lu, Y., Wang, H., Shi, J., Lu, H., Nielsen, J., 2025. Computational biology predicts metabolic engineering targets for increased production of 103 valuable chemicals in yeast. Proc Natl Acad Sci U S A. 122**,** e2417322122.

Ebrahim, A., Lerman, J. A., Palsson, B. O., Hyduke, D. R., 2013. COBRApy: COnstraints-Based Reconstruction and Analysis for Python. BMC Syst Biol. 7**,** 74.

Gurobi Optimization, L. L. C., Gurobi Optimizer Reference Manual.

Holland, J. H., 1992. Genetic Algorithms. Scientific American. 267**,** 66-72.

Schneider, P., Bekiaris, P. S., von Kamp, A., Klamt, S., 2022. StrainDesign: a comprehensive Python package for computational design of metabolic networks. Bioinformatics. 38**,** 4981-4983.

Wang, H., Zhang, M., Zhang, C., He, S., Liao, W., Zhu, R., Zhou, Y. J., Lu, H., 2025. StrainOptimizer empowers rational cell factory design through multi-scale metabolic models with expression and proteome constraints. bioRxiv.

Zhang, J., Sanderson, A. C., 2009. JADE: Adaptive Differential Evolution With Optional External Archive. IEEE Transactions on Evolutionary Computation. 13**,** 945-958.
